# Repurposed COMT Inhibitors Tolcapone and Entacapone Selectively Suppress Aggregation and Seeding of P301 Mutant TAU in Human Neuronal Models

**DOI:** 10.64898/2026.04.20.719548

**Authors:** Ihor Kozlov, Yu-Sheng Hung, Sudeep Roy, Alladi Charanraj Goud, Roman Kouřil, Yu-Hui Wong, Viswanath Das

## Abstract

**Background and Purpose:** Pathogenic aggregation and propagation of seed-competent TAU assemblies drive tauopathies. *MAPT* P301 mutations accelerate aggregation and enhance seed competence, yet pharmacological strategies selectively targeting these pathogenic species remain limited. We investigated whether the clinically approved catechol-O-methyltransferase inhibitors tolcapone (TOL) and entacapone (ENT) preferentially modulate mutant TAU aggregation and seeding.

**Experimental Approach:** TOL and ENT effects on TAU aggregation were evaluated via cell-free assays, surface plasmon resonance (SPR), and *in silico* docking. Functional consequences of compound-modified fibrils were assessed in mutant TAU-expressing SH-SY5Y cells. Translational relevance was examined in human induced pluripotent stem cell (hiPSC)-derived neurons exposed to pathogenic K18 fibrils, followed by post-seeding compound treatment.

**Key Results:** Both compounds dose-dependently inhibited TAU aggregation, exhibiting greater potency, stronger SPR binding affinities, and more favorable computed interaction energies for P301S mutant versus wild-type TAU. Fibrils formed with TOL or ENT induced less downstream TAU oligomerization and phosphorylation in SH-SY5Y cells, with TOL showing superior protection. In hiPSC-derived neurons, post-seeding treatment with either compound decreased fibril-induced, sarkosyl-insoluble TAU aggregation and phosphorylation without overt cytotoxicity.

**Conclusion and Implications:** TOL and ENT preferentially inhibit the aggregation and seeding of pathogenic P301 mutant TAU. This supports mutation-focused pharmacological strategies and highlights catechol scaffolds as viable starting points for the development of disease-modifying therapeutics. Future research must determine the precise interaction mechanisms with aggregation intermediates and evaluate *in vivo* efficacy in animal models.

## INTRODUCTION

Intracellular aggregation of the microtubule-associated protein TAU defines tauopathies (Strang et al., 2019). This pathology emerges early, spreading progressively through neuronal networks in a prion-like manner, driving disease progression and clinical heterogeneity (Cope et al., 2018; Kaufman et al., 2018; Kim et al., 2020; Vogel et al., 2020; Franzmeier et al., 2021). Because TAU pathology often precedes substantial neuronal loss and brain atrophy, pathogenic TAU species are considered upstream drivers of neurodegeneration rather than passive by-products (Xia et al., 2017; La Joie et al., 2020).

Specific TAU assemblies act as seeds, templating the conversion of soluble TAU into aggregation-prone conformers to enable cell-to-cell spread (Brunello et al., 2019). Unlike the low-level aggregation seen in normal aging, tauopathies feature the accumulation of stable, seed-competent TAU species that propagate efficiently (Vogel et al., 2020). This seeding and propagation are influenced by neuronal activity, cellular stress, and *MAPT* gene mutations (Strang et al., 2019).

TAU aggregation is primarily driven by its microtubule-binding domain, specifically the amyloidogenic motifs VQIINK (PHF6*) and VQIVYK (PHF6), which are central to fibril formation and seeding (Li and Lee, 2006; Ganguly et al., 2015; Stöhr et al., 2017). In wild-type (WT) TAU, local secondary structure partially shields VQIVYK, limiting spontaneous aggregation (Mirbaha et al., 2018). Disrupting this structure via post-translational modifications or mutations exposes these sequences, enhancing aggregation (Chen et al., 2019).

Mutations at residue P301 (e.g., P301L, P301S) are highly pathogenic. They destabilize the R2–R3 interface, weakening interactions that restrain β-sheet formation, thereby accelerating aggregation and seed formation (Fischer et al., 2009; Chen et al., 2019). P301-mutant assemblies show greater stability, seeding activity, and clearance resistance than WT TAU. Accordingly, cryo-electron microscopy of tauopathy filaments reveals VQIVYK as a core structural element in pathogenic fibrils (Fitzpatrick et al., 2017; Falcon et al., 2018).

Many aggregation inhibitors (small molecules, peptides, antibodies) that succeed *in vitro* fail to block TAU seeding and propagation in cellular or *in vivo* systems (Soeda et al., 2019; Soeda and Takashima, 2020; Hill et al., 2022). This highlights the need to specifically suppress seed-competent TAU species rather than just general aggregation. Structure-based inhibitors targeting motifs like VQIVYK show promise in blocking seeding, even against patient-derived aggregates (Zheng et al., 2011; Seidler et al., 2018, 2019).

Catechol-containing compounds, including clinically approved catechol-O-methyltransferase (COMT) inhibitors like tolcapone (TOL) and entacapone (ENT), interfere with TAU aggregation. Early cell-free studies showed they inhibit PHF6 hexapeptide aggregation, identifying the nitrocatechol moiety as a key pharmacophore (Mohamed et al., 2013). Later work found catechols suppress TAU oligomerization via cysteine-dependent interactions and reduce pathological accumulation in models (Soeda et al., 2015). However, these studies focused on bulk aggregation or early oligomers, leaving it unclear if these compounds modulate seed-competent TAU assemblies or distinguish between WT and mutant conformers.

Here, we investigate whether TOL and ENT preferentially suppress the aggregation and seeding of pathogenic P301 TAU. Using biochemical assays, SH-SY5Y cells, and human induced pluripotent stem cell–derived neurons expressing P301L TAU, we assess their effects on aggregation kinetics, seed competence, and intracellular propagation. Our findings highlight clinically approved small molecules that selectively target pathogenic P301 assemblies, establishing a mechanistic framework for mutation-focused strategies against seed-competent TAU in tauopathies.

## METHODS

### Materials and Reagents

TAU peptides corresponding to WT-R2R3 and P301S-R2R3 TAU constructs were synthesized and handled as described previously (Goud et al., 2025). Unless otherwise mentioned, all reagents were purchased from Sigma-Aldrich (St. Louis, MO, USA). TOL (Cat. # SML0150) and ENT (Cat. # SML0654) were dissolved in HPLC-grade DMSO and stored at -80°C. All intermediate drug dilutions were prepared in PBS (pH 7.4) supplemented with 0.05% Tween-20 and 5% DMSO. Unless otherwise indicated, compounds were added in a volume-normalized manner such that the final DMSO concentration did not exceed 0.125% (v/v) in any condition, including vehicle controls.

### *In Vitro* TAU Aggregation Assay

TAU aggregation kinetics were monitored using ThT fluorescence assays as described previously (Goud et al., 2025). TAU peptides stored at −80 °C were thawed on ice and centrifuged at 13,000 rpm for 45 s before use. Aggregation reactions were prepared in aggregation buffer (20 mM Tris-HCl, pH 7.4, 100 mM NaCl, 1 mM EDTA) containing 15 µM ThT (Sigma-Aldrich, Cat. #T3516-5G) and 50 µM TAU peptide, in the absence or presence of TOL or ENT at concentrations ranging from 0.10-100 µM. Final reaction volume was 40 µL per well in clear-bottom, black 384-well plates sealed with TopSealA-PLUS to prevent evaporation.

Plates were incubated at 37 °C with constant orbital agitation (1 mm amplitude, 432–1000 rpm depending on the reader). ThT fluorescence (λex 485 nm, λem 535 nm) was recorded every 5–10 min for up to 30-35 h using Infinite Pro 200 (Tecan).

Aggregation kinetics were analyzed using GraphPad Prism (version 10, San Diego, CA, USA). ThT fluorescence-time profiles were used to calculate the area under the curve (AUC) for each condition as an integrated measure of aggregation. AUC values were normalized to the untreated control (CTL). Concentration–response curves were generated by plotting normalized AUC values against compound concentration and fitted using three-parameter logistic models to derive half-maximal inhibitory concentration (IC_50_) values

### Transmission Electron Microscopy

Aggregation end-point samples were analyzed by transmission electron microscopy to assess fibril morphology. Samples of WT and P301S R2R3 TAU constructs aggregated in the absence or presence of TOL or ENT were prepared for imaging after 48 h of incubation.

Aliquots (2–3 µL) were applied to glow-discharged carbon-coated copper grids and incubated for 1.5 min. Excess liquid was removed with filter paper, and grids were air-dried prior to imaging.

Grids were imaged using a Tecnai G2 F20 transmission electron microscope (FEI Technologies, Hillsboro, USA) equipped with an Eagle 4K CCD camera. Approximately 15 images were acquired per sample from multiple grid regions. Fibril morphology was assessed qualitatively, and fibril parameters were analysed using NIH ImageJ software (RRID: SCR_003070).

Fibril width was quantified by measuring the edge-to-edge distance perpendicular to the fibril axis using an ImageJ-based line tool and macro-assisted workflow. Images with a standardized scale were analyzed by manually drawing two lines, each with an identical number of points, along opposing fibril edges, oriented parallel to the fibril axis. The macro automatically generated perpendicular distance measurements between corresponding points and calculated an average fibril width per measurement. Multiple fibrils were measured per image, pooled per condition for statistical analysis, and the ImageJ macro used for this analysis is provided as supplementary code via Zenodo (Data Availability Statement).

### Generation and Processing of TAU Aggregates for Cell-Based Seeding Assays

TAU aggregation reactions for cell seeding assays were performed in aggregation buffer without ThT, and as described previously (Goud et al., 2025). Briefly, aggregation reactions were allowed to proceed for 30–35 h, after which reaction products were pooled, and the TAU concentration was determined using a BCA protein assay. Aggregation mixtures were then standardized to a final concentration of 100 nM TAU peptide, sonicated using a Branson Ultrasonic™ Sonifier Cup Horns (Marshall Scientific, Hampton, NH, USA) at 30% amplitude, 15 s ON/15 s OFF cycles for 1 min, and used immediately for cell transduction experiments.

### Cell lines and Human Induced Pluripotent Stem Cells

SH-SY5Y neuroblastoma cells, originally obtained from ATCC (Manassas, VA, USA; Cat. No. CRL-2266; RRID: CVCL_0019), were engineered to stably express a doxycycline-inducible AAVS1-integrated TAUP301L-2N4R-EGFP (Addgene plasmid #132,393; http://n2t.net/addgene:132393; RRID: Addgene_132393) as described previously (Devyatov et al., 2025). Stable integration was verified by puromycin antibiotic selection with limiting dilution cloning, followed by confirmation of doxycycline-inducible EGFP expression using fluorescence microscopy and Western blot analysis (Devyatov et al., 2025).

SH-SY5Y cells were maintained in Dulbecco’s Modified Eagle’s Medium high glucose (4.5 g/L) with L-glutamine (Lonza, #12-604F), supplemented with 10% fetal calf serum and 1% penicillin–streptomycin, in a humidified incubator at 37 °C with 5% CO_2_. Cells were maintained at approximately 80% confluency and passaged every 3–4 days and used between passages 3 and 15 for all experiments after thawing. Cell lines were authenticated periodically by short tandem repeat (STR) profiling and tested for mycoplasma contamination bi-weekly according to established laboratory protocols (2022).

The human induced pluripotent stem cell (hiPSC) line KOLF2.1J was obtained from The Jackson Laboratory (#JIPSC1000). hiPSCs were maintained in StemFlex medium (Invitrogen, #A3349401) on culture dishes precoated with recombinant human vitronectin (5 µg/mL; Invitrogen, #A14700). Cells were passaged every 4–5 days using 0.5 mM EDTA and reseeded at a ratio of 1:5–1:10. Culture medium was refreshed according to the manufacturer’s instructions. hiPSCs were routinely monitored for morphology and growth characteristics and confirmed to be mycoplasma-negative using Myco-Detect Fast qPCR Kits (Topgen Biotech., #QM050) prior to downstream differentiation experiments; cells were used between passages 5–15 after thawing.

### SY5Y-TAUP301L Cell Differentiation, and Seeding

For neuronal differentiation, SY5Y-TAUP301L cells were transferred to Neurobasal™-A medium supplemented with L-glutamine, B-27, penicillin-streptomycin, retinoic acid, and recombinant human BDNF, as described previously (Devyatov et al., 2025). Expression of TAUP301L-EGFP was induced by adding doxycycline (0.5 µg/mL) 3 days before seeding and maintained throughout the experiment.

For seeding assays, differentiated SY5Y-TAUP301L cells were transduced with TAU aggregation products at a final concentration of 100 nM using TurboFect™ transfection reagent in Opti-MEM™, as described previously (Devyatov et al., 2025; Goud et al., 2025). Transfection mixtures were incubated for 15 min at room temperature prior to addition to cells. Cells were incubated for 72 h after transduction, then processed for Triton X-100 TAU fractionation and Western blot analysis as described below.

### Generation of Isogenic iPSC Lines Using CRISPR/Cas9

Guide RNA (gRNA) sequences targeting *MAPT* exon 10 were designed using the TrueGuide™ CRISPR gRNA Design Tool (Thermo Fisher Scientific). The selected gRNA (CRISPR1107020_SGM; sequence: AATATCAAACACGTCCCGGG) was synthesized and assembled with Cas9 nuclease (Invitrogen) to form Cas9 ribonucleoprotein complexes (Cas9 RNPs). Cas9 RNPs were electroporated into hiPSCs using the Neon™ system (Invitrogen). For P301L editing by homology-directed repair, Cas9 RNPs were co-electroporated with a piggyBac donor plasmid carrying the P301L mutation using the following Neon settings: 1,400 V, 10 ms, 3 pulses (program #22). Cells were treated with puromycin (1 mg/L) for 48 h starting on day 3 post-electroporation, followed by a second 48-h puromycin selection on day 10. Individual surviving colonies were manually picked two weeks after transfection, expanded, and screened by PCR. Correctly edited clones were verified by Sanger sequencing (NYCU Sequencing Services). For downstream neuronal experiments in this study, one validated MAPT(P301L) isogenic clone was used, and all treatment comparisons (vehicle vs. TOL/ENT) were performed within the same genetic background.

### Generation of Induced Neurons (iNs) from hiPSCs

hiPSCs were transfected with PB-TO-hNGN2 (Addgene plasmid #172115; a gift from iNDI & Michael Ward) and cultured in StemFlex medium containing puromycin (1 mg/L) for one week.

For differentiation, hiPSCs were dissociated with 0.5 mM EDTA and replated onto vitronectin-coated dishes in Essential 8 medium (Invitrogen, #A1517001) supplemented with Y-27632 (5 µM) (day −1). On day 0, the medium was replaced with DMEM/F12 supplemented with N2, NEAA, BDNF (10 µg/L), NT-3 (10 µg/L), mouse laminin (0.1 mg/L), and doxycycline (2 mg/L). On day 2, cells were dissociated and replated onto poly-D-lysine/laminin-coated plates at 6–8 × 10^4^ cells/cm^2^ in Neurobasal medium containing B27, GlutaMAX, mouse laminin, BDNF, and NT-3. Ara-C (2.5 µM) was added on day 5 for 48 h to inhibit the proliferation of undifferentiated cells. Afterward, half of the medium was replaced every 3 days with fresh Neurobasal medium supplemented with B27/GlutaMAX/laminin/BDNF/NT-3.

### Preparation and Transfection of sK18f

Pre-formed P301L mutant human TAU (K18) fibrils were purchased from GeneTex (GTX17675-pro). Fibrils were diluted to 0.2 µg/µL in phosphate-buffered saline (PBS; 137 mM NaCl, 3 mM KCl, 7 mM Na□HPO□, 1.5 mM KH□PO□, pH 7.4) and sonicated on ice for 10 cycles of 30-s pulses with 30-s rest at 30–40% amplitude. Sonicated K18 fibrils (sK18f) were aliquoted and stored at −80°C.

Neuronal transfection was performed using Xfect (Takara, #631318) according to the manufacturer’s instructions. For 24-well plates, each well received 0.3 µg sK18f and 1.5 µL Xfect; for 35-mm dishes, 1.5 µg sK18f and 7.5 µL Xfect were applied.

### Compound Treatment and ER Stress Induction

One day after sK18f exposure, the entire culture medium was removed and replaced with Neurobasal medium supplemented with B27, GlutaMAX, mouse laminin, BDNF, and NT-3, containing TOL or ENT (2, 10, or 50 µM). Media and compounds were refreshed every 3 days and maintained for three weeks. To model a stress-sensitized neuronal environment relevant to tauopathy, endoplasmic reticulum stress was induced by applying 0.4 µM thapsigargin for one week during week 6 of differentiation (Devina et al., 2022).

### Fluorescent Labeling of sK18f

For fluorescent labeling, 100 µg sK18f was buffer-exchanged into 0.1 M sodium bicarbonate (pH 8.3) using an Amicon Ultra-0.5 centrifugal filter (Merck, #UFC501024) and concentrated to 45 µL. Alexa Fluor 647 NHS ester (100 µg in 10 µL DMSO; Invitrogen, #A37573) was added (5 µL) and incubated at 25°C with shaking at 1,500 rpm on a ThermoMixer C. After labeling, the reaction was washed four times with 500 µL PBS using an Amicon Ultra-0.5 filter to remove unreacted dye.

### TAU Fractionation

#### Triton X-100 fractionation of SY5Y-TAUP301L cells

Triton X-100-soluble and -insoluble fractions from SY5Y-TAUP301L cells were prepared as described previously (Devyatov et al., 2025), with minor modifications. Briefly, cells were lysed in 1× TBS containing 0.05% Triton X-100, supplemented with protease and phosphatase inhibitors. Lysates were first centrifuged at 1,000 × g for 10 min at 4°C to remove cell debris. The resulting supernatant was then ultracentrifuged at 24,400 × g for 1 h at 4°C. The supernatant from this step was collected and stored as the Triton X-100–soluble fraction. The remaining pellet was washed once with 1× TBS containing 0.05% Triton X-100, supplemented with protease and phosphatase inhibitors, by centrifugation at 24,400 × g for 3 min at 4°C. The supernatant was discarded, and the final pellet was resuspended in RIPA buffer (supplemented with protease and phosphatase inhibitors) at the same volume as the corresponding soluble fraction and used as the Triton X-100–insoluble fraction. Protein concentration was determined in the soluble fraction using the BCA assay.

#### Sarkosyl fractionation of hiPSC-derived neurons

After treatment with sK18f with or without TOL or ENT, hiPSC-derived neurons were lysed in high-salt lysis buffer containing 10 mM HEPES (pH 7.5), 800 mM NaCl, 1 mM EGTA, 10% sucrose, protease inhibitor cocktail (Roche, #4693116001), phosphatase inhibitor cocktail I (TargetMol, #C0002), and phosphatase inhibitor cocktail II (TargetMol, #C0003). Cells were homogenized by pipetting, and 1/30 of the volume was added to 30% sarkosyl (Sigma, #61747) to reach a final concentration of 1%. Lysates were incubated at room temperature with gentle inversion for 1 h and centrifuged at 15,000 rpm for 1 h at 4°C. The supernatant was collected as the sarkosyl-soluble fraction. The pellet was washed once with PBS and resuspended in pellet buffer containing 10 mM HEPES (pH 7.5), 2 mM MgCl□, 1% Triton X-100, 1 U/µL benzonase (Millipore, #E1014), protease inhibitor cocktail, and phosphatase inhibitor cocktail I/II. The pellet was disrupted by pipetting, incubated at 37°C for 30 min to digest genomic DNA and RNA, and then stored at –20°C as the sarkosyl-insoluble fraction.

### Western Blotting

For hiPSC-derived neurons, sarkosyl-soluble and sarkosyl-insoluble fractions prepared as described above were used directly for immunoblotting. For SY5Y-TAUP301L cells, Triton X-100–soluble and –insoluble fractions were prepared as described above, and the Triton X-100-insoluble fraction was used for the detection of aggregated and phosphorylated TAU species, while the corresponding soluble fraction was used for GAPDH detection as a loading control.

Protein samples were mixed with 4× SDS sample buffer (250 mM Tris-HCl pH 8.5, 8% SDS, 40% glycerol, 2 mM EDTA, 0.075% Serva Blue G250, 0.025% phenol red, and 10% 2-mercaptoethanol) and denatured at 75°C for 15 min. Denatured samples were separated on 8% SDS–PAGE gels at 70 V for 20 min, followed by 120 V for 1 h, and transferred to 0.45 µm PVDF membranes (Millipore) in transfer buffer (25 mM Tris, 192 mM glycine, 20% methanol) at 70 V for 2.5 h.

Membranes were blocked in 1× TBS containing 0.05% Tween-20 (TBST) and either 5% skimmed milk or 5% bovine serum albumin for 30-60 min at room temperature and incubated with primary antibodies (Table 1) overnight at 4°C. After three 10-minute washes in 1× TBST, membranes were incubated with HRP-conjugated secondary antibodies or Alexa Fluor 488 488-conjugated secondary antibodies for 2-4 h at 4°C, washed three times. For reprobing, membranes were stripped with stripping buffer (1.5% glycine, pH 2.2, 1% Triton X-100, 0.1% SDS) or quenched with 3% H_2_O_2_ in 1× TBST for 30 min to remove residual antibodies or HRP activity, respectively, followed by two washes with TBST before the next antibody incubation.

**Table 1.** List of antibodies used in the study.

| Antibody | Host species | Dilution | Manufacturer | Cat. #; RRID |
| --- | --- | --- | --- | --- |
| TAU-5 | Mouse | 1:500 and 1:1000 (WB) | Thermo Fisher Scientific | AHB0042; RRID: AB_2536235 |
| pTAU (Ser262) | Rabbit | 1:1000 | Thermo Fisher Scientific | OPA1-03142; RRID: AB_325643 |
| Anti-TAU (T22), Oligomeric | Rabbit | 1:1000 | Millipore | ABN454; RRID: AB_2888681 |
| GAPDH | Rabbit | 1:4000 | Cell Signaling Technologies | 2118S; RRID: AB_561053 |
| pTAU (S202/T205) [AT8] | Mouse | 1:500 (WB); 1:200 (IF) | Addgene | 216150; RRID: AB_3094509 |
| pTAU (S262) | Rabbit | 1:500 | Santa Cruz Biotechnology | sc-32828; RRID: AB_1129983 |
| TAU | Rabbit | 1:4000 | GeneTex | GTX639564; RRID: AB_3738550 |
| GAPDH | Mouse | 1:10000 | GeneTex | GTX627408; RRID: AB_11174761 |
| TUJ1 | Rabbit | 1:1000 | GeneTex | GTX637307; RRID: |
|  |  |  |  | AB_3738549 |
| Goat anti-mouse IgG-HRP | Goat | 1:10000 | Bethyl laboratories | A90-216P;<br>RRID:<br>AB_67184 |
| Goat anti-rabbit IgG-HRP | Goat | 1:10000 | Bethyl laboratories | A120-201P;<br>RRID:<br>AB_10631424 |
| F(ab')<sub>2</sub>-Goat anti-Rabbit IgG (H +L) Cross-Adsorbed Secondary Antibody, Alexa Fluor™ 488 | Goat | 1:2000 | Thermo Fisher Scientific | A-11070;<br>RRID:<br>AB_2534114 |
| Donkey anti-Mouse IgG (H +L) Highly Cross-Adsorbed Secondary Antibody, Alexa Fluor™ 488 | Donkey | 1:2000 | Thermo Fisher Scientific | A-21202;<br>RRID:<br>AB_141607 |
| F(ab')<sub>2</sub>-Goat anti-Rabbit IgG (H+L) Cross-AdsorbedSecondary Antibody, Alexa Fluor Plus 647 | Goat | 1:500 | Invitrogen | A48285;<br>RRID:<br>AB_2896349 |
| Goat anti-Mouse IgG (H+L) Highly Cross-Adsorbed Secondary Antibody, Alexa Fluor™ 488 | Goat | 1:400 | Invitrogen | A-11029;<br>RRID:<br>AB_2534088 |

For HRP-based detection, membranes were developed using Immobilon Western HRP substrate (Millipore), and chemiluminescent signals were detected using an ImageQuant LAS 4000 imaging system. For fluorescence-based detection, Alexa Fluor 488 signals were detected using a ChemiDoc MP imaging system (Bio-Rad). All densitometric analyses were performed using ImageJ.

### Immunofluorescence Staining

Cells were fixed with 4% paraformaldehyde in PBS for 20 min at room temperature, rinsed three times with PBS, and permeabilized with 0.1% Triton X-100 in PBS for 5 min. Samples were blocked in PBS containing 3% bovine serum albumin (Sigma) and 1% goat serum (Gibco) for 1 h at room temperature, followed by incubation with primary antibodies (Table 1) overnight at 4°C. After three washes with PBS, appropriate secondary antibodies were applied for 1 h at room temperature.

ProteoStat staining was then performed using the ProteoStat aggregation detection kit (Enzo Life Sciences, ENZ-51035) according to the manufacturer’s instructions. After final washes, samples were mounted using VECTASHIELD® Antifade Mounting Medium (Vector Laboratories, H-1000). Images were acquired using either a Zeiss LSM900 confocal microscope equipped with Airyscan 2 and an AxioCam 702 MONO camera, or a Zeiss fluorescence microscope equipped with an Andor Zyla CMOS camera, and processed using ImageJ.

### LDH Release Assay

Lactate dehydrogenase (LDH) release was quantified using the CytoTox 96® Non-Radioactive Cytotoxicity Assay Kit (Cat. #G1780, Promega) according to the manufacturer’s instructions. After 4 weeks of differentiation, iPSC-derived neurons were treated with or without sK18f for 1 hour, followed by administration of VQIVYK inhibitors. DMSO (0.25%, v/v) was used as the vehicle control. Neurons were subsequently maintained in culture with VQIVYK inhibitors for 5 days. To generate the 100% LDH release control, Triton X-100 was added to untreated neurons at a final concentration of 0.8% (v/v) and incubated at 37°C for 45 minutes. Culture supernatants were then collected and transferred to 96-well plates. LDH assay reagent was added according to the manufacturer’s protocol and incubated for 30 minutes at room temperature. The reaction was terminated by adding stop solution, and absorbance was measured at 490 nm using a TECAN Infinite 200 PRO microplate reader.

The percentage of cell death was calculated using the following formula:

Cell death (%) = (Sample LDH release – Blank)/(Maximum LDH release – Blank) × 100

### Live/Dead Cell Viability Assay

Cell viability was assessed based on intracellular esterase activity and plasma membrane integrity using the LIVE/DEAD® Viability/Cytotoxicity Kit for mammalian cells (Cat. #L3224, Invitrogen), following the manufacturer’s instructions. After 4 weeks of differentiation and maturation, iPSC-derived neurons were treated with sK18f for 1 hour, followed by incubation with VQIVYK inhibitors or DMSO vehicle control for 5 days. Cells were then stained with 0.2 µM calcein AM and 0.2 µM ethidium homodimer-1 at 37°C for 10 minutes. Fluorescence images were acquired using an Olympus IX-83 fluorescence microscope. Live and dead cells were quantified using Fiji (ImageJ) software.

### Statistical Analysis

All data were analyzed using GraphPad Prism. Data are presented as mean ± SEM. Statistical significance was defined as P < 0.05. The statistical tests used for each experiment are specified in the corresponding figure legends.

## RESULTS

### TOL and ENT suppress TAU aggregation kinetics with enhanced effects on mutant TAU

The effects of TOL and ENT on TAU aggregation were monitored using a thioflavin T (ThT) binding assay under conditions supporting spontaneous fibril formation, using WT R2R3 TAU (WT-R2R3) or P301S-mutant R2R3 TAU (P301S-R2R3) constructs. These peptide constructs represent a minimal aggregation-competent TAU model described previously (Das et al., 2025). Both TOL and ENT reduced aggregation of WT-R2R3 and P301S-R2R3 compared with untreated controls (Fig. 1B, C). TEM of aggregation end-point samples showed abundant fibrillar assemblies under control conditions, whereas reactions performed in the presence of TOL or ENT contained markedly fewer and morphologically altered fibrillar structures (Fig. 1D). Quantitative analysis showed that fibrils formed in the presence of either compound exhibited a shift toward reduced width distributions for both WT-R2R3f and P301S-R2R3f relative to control fibrils (Fig. 1E, F).

**Figure 1.**
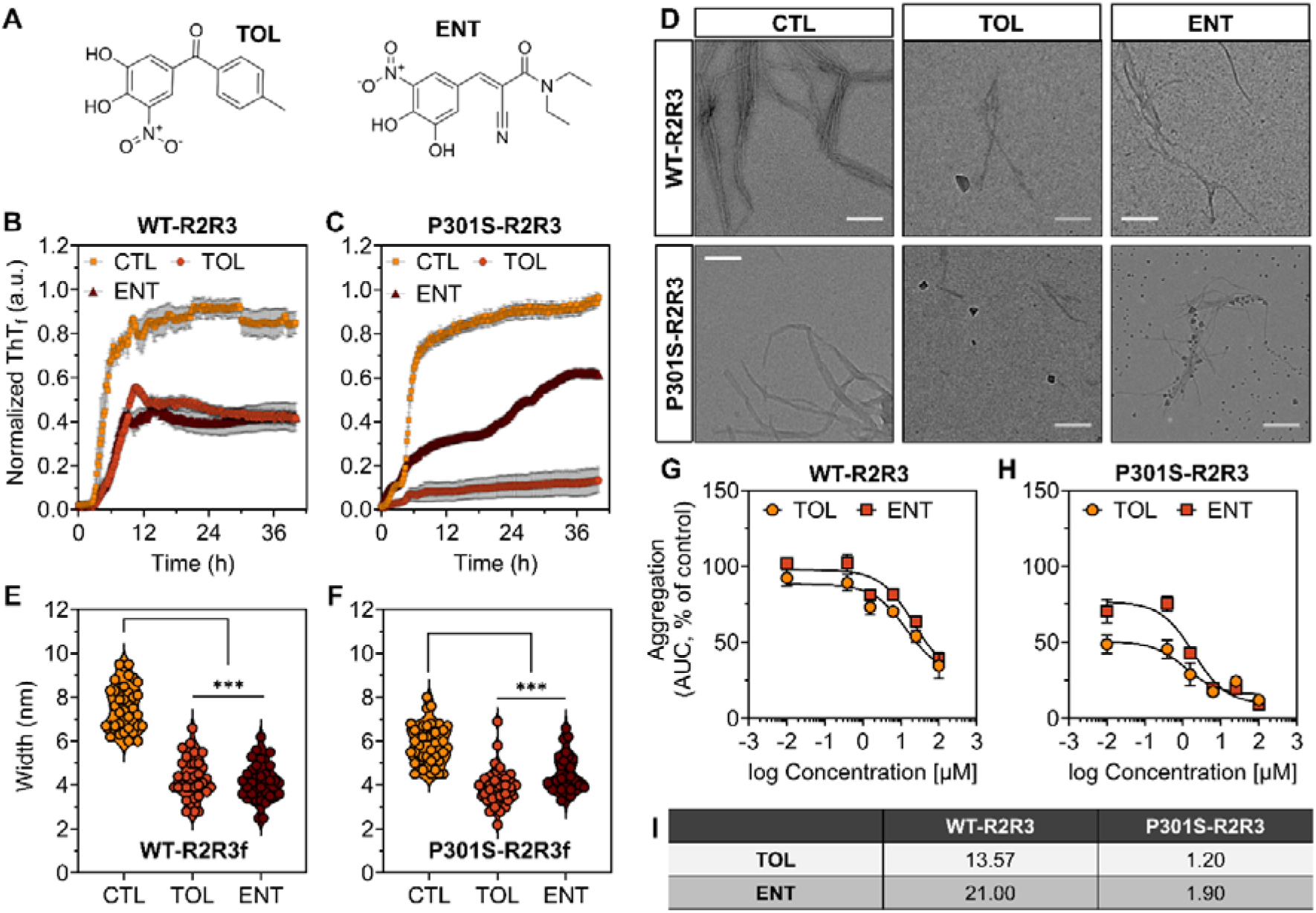
TOL and ENT preferentially inhibit mutant TAU aggregation. (**A**) Chemical structures of TOL and ENT. (**B, C**) ThT aggregation kinetics of WT-R2R3 (B) and P301S-R2R3 (C) TAU constructs incubated under standard aggregation conditions in the absence (CTL) or presence of 10 µM TOL or ENT. Curves represent mean ± SEM (*n* = 3 independent experiments). (**D**) TEM images of aggregates formed from WT-R2R3 and P301S-R2R3 TAU constructs under CTL conditions or in the presence of TOL or ENT. Scale bar, 100 nm. (**E, F**) Violin plots showing the distribution of fibril width measurements for WT-R2R3f (E) and P301S-R2R3f (F) formed under CTL conditions or in the presence of TOL or ENT. Each violin represents pooled measurements from multiple fibrils across 2–3 independent TEM images per condition; central lines indicate the median and interquartile range. ***P < 0.001, one-way ANOVA with Dunnett’s multiple comparisons test. (**G, H**) Concentration-dependent inhibition of WT-R2R3 (G) and P301S-R2R3 (H) TAU aggregation by TOL and ENT, expressed as AUC of ThT fluorescence normalized to the corresponding untreated control (aggregation, % of control). Data are mean ± SEM (*n* = 3 independent experiments). Corresponding ThT kinetic traces used for AUC calculations are shown in Fig. S1. (I) IC_50_ (µM) values for inhibition of TAU aggregation by TOL and ENT, derived from AUC-based concentration–response analyses of ThT fluorescence kinetics shown in panels G and H.

To compare inhibitory efficacy across constructs and compounds, aggregation assays were performed over a range of TOL and ENT concentrations. Aggregation was quantified by calculating the area under the curve (AUC) of ThT fluorescence kinetics for each condition and normalizing values to the corresponding untreated control, as described previously (Goud et al., 2025). Normalized aggregation decreased in a concentration-dependent manner for both WT-R2R3 and P301S-R2R3 in response to TOL or ENT (Fig. 1G, H). Concentration–response relationships were fitted using three-parameter logistic models to derive half-maximal inhibitory concentration (IC_50_) values for aggregation inhibition (Fig. S1). These analyses revealed consistently lower IC_50_ values for P301S-R2R3 compared with WT-R2R3 for both compounds (Fig. 1I).

### TOL and ENT bind P301S TAU more strongly than WT TAU

To test whether the preferential functional effects of TOL and ENT on mutant TAU reflect differential compound–TAU interactions, we quantified their binding to monomeric WT-R2R3 and P301S-R2R3 TAU constructs using surface plasmon resonance (SPR). Binding was concentration-dependent for both compounds across WT-R2R3 and P301S-R2R3 TAU, and global kinetic fitting yielded equilibrium dissociation constants (KD) together with association (Kon) and dissociation (Koff) rate constants (Table 2; Fig. S2). TOL bound WT-R2R3 TAU with a higher KD than P301S-R2R3 TAU, indicating stronger binding to the mutant. ENT displayed overall weaker binding than TOL but maintained a lower KD for P301S-R2R3 than WT-R2R3.

**Table 2.** SPR kinetic parameters for TOL and ENT. KD, Kon, and Koff values for TOL and ENT binding to WT-R2R3 and P301S-R2R3 TAU constructs, determined by surface plasmon resonance. Corresponding sensorgrams are shown in Fig. S4. Lower KD values indicate higher apparent binding affinity.

|  | **R2R3** | **KD (M)** | **Kon (M<sup>-1</sup>s<sup>-1</sup>)** | **Koff (s<sup>-1</sup>)** |
| --- | --- | --- | --- | --- |
| **TOL** | WT | $1.50 \times 10^{-7}$ | 53.24 | $8.01 \times 10^{-6}$ |
| | P301S | $3.30 \times 10^{-8}$ | 45.17 | $1.49 \times 10^{-6}$ |
| **ENT** | WT | $2.11 \times 10^{-4}$ | 7.431 | $1.57 \times 10^{-4}$ |
| | P301S | $1.25 \times 10^{-5}$ | 12.33 | $1.54 \times 10^{-4}$ |

To provide structural context, docking was performed using AlphaFold2-derived models of WT and P301S TAU (Fig. 2; Fig. S3). All structural models were evaluated for confidence and stereochemical quality prior to docking (Fig. S4–S8). Docking identified defined binding regions for both ENT and TOL in WT and P301S TAU (Fig. 2), with predicted binding scores in a comparable range across variants (Table S1). Consistent with the SPR measurements, predicted binding energies were modestly more favorable for P301S than for WT (Table S1). The additional model, P301L, included for comparative analysis, showed a similar binding pattern and energy range (Fig. S8; Table S1). Molecular dynamics–based binding analyses (Tables S2–S7) showed average MM-GBSA binding energies across ten replicates that supported more favorable predicted interactions of ENT and TOL with mutant TAU (P301S and P301L) than with WT TAU.

**Figure 2.**
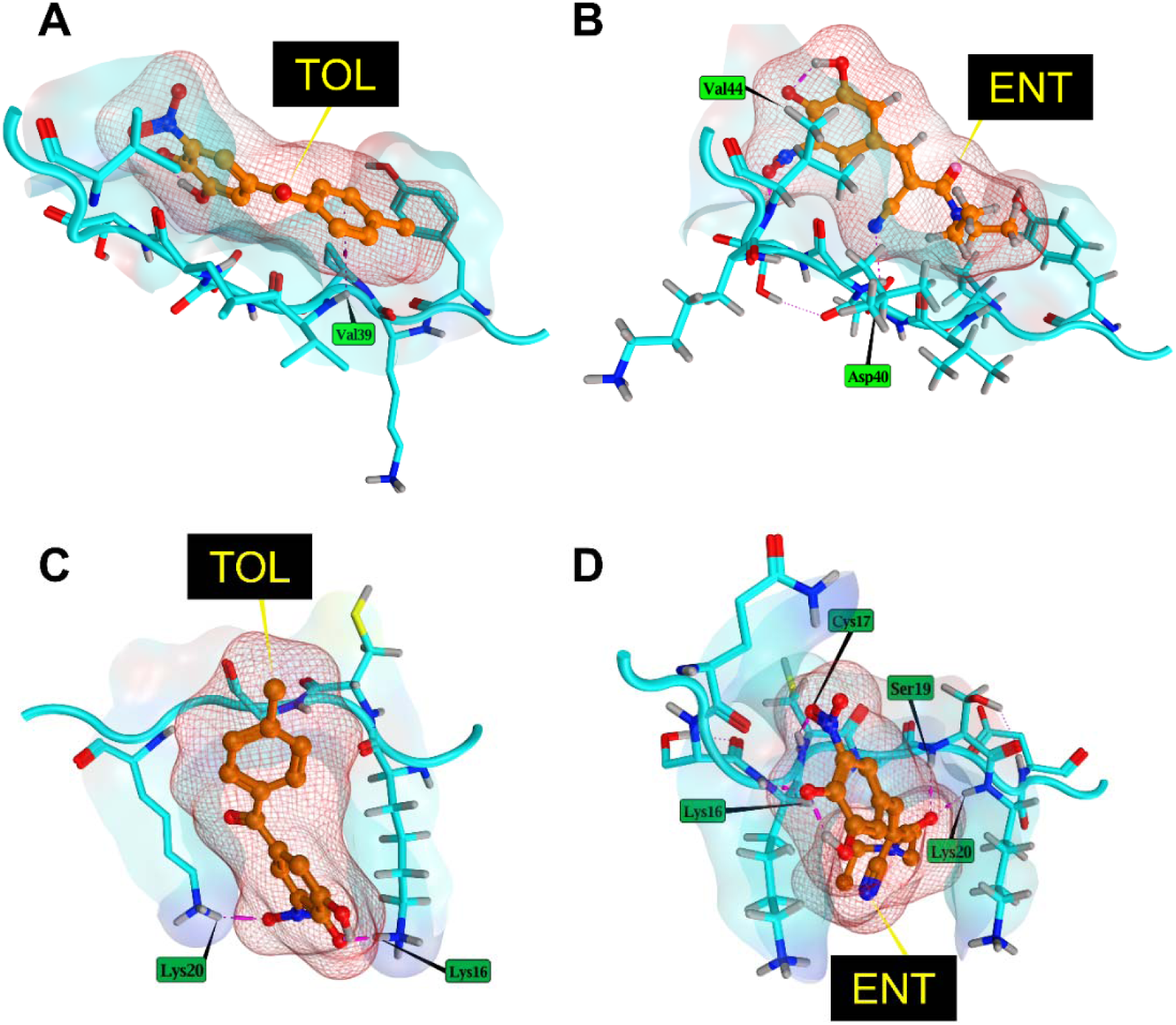
Docking interactions of TOL and ENT with WT and P301S tau filament models. Predicted binding poses of TOL (A, C) and ENT (B, D) docked to AlphaFold2-derived WT tau filaments (A, B) and P301S tau filaments (C, D). The tau filament is shown as a ribbon in surface representation, with selected interacting residues displayed as sticks. Ligands are represented as space-filling models colored by atom type. Docking was performed using MOE 2024.01. Interactions are depicted as hydrogen bonds (yellow dashed lines), hydrophobic contacts (pink), and potential ionic interactions (violet lines).

### R2R3f formed in the presence of TOL or ENT show reduced pathogenicity in neuron-like SY5Y cells expressing TAUP301L-2N4R

To assess whether compound-modified TAU assemblies differ in their cellular pathogenicity, we examined their effects in SH-SY5Y cells expressing TAUP301L-2N4R-EGFP (SY5Y-TAUP301L). Differentiated cells were transduced with WT-R2R3f or P301S-R2R3f, formed either in the absence of compounds (control, CTL) or in the presence of TOL or ENT during fibril assembly, as described previously (Goud et al., 2025).

Immunoblot analysis of Triton X-100–insoluble fractions showed that seeding with WT-R2R3f or P301S-R2R3f generated under CTL conditions markedly increased levels of T22-reactive oligomeric and Ser262-phosphorylated TAU compared with buffer-only CTL (Fig. 3A). In contrast, cells transduced with fibrils formed in the presence of TOL or ENT exhibited substantially reduced levels of both T22-positive TAU species and pTAU (Ser262) for both WT-R2R3f and P301S-R2R3f. Total TAU in the Triton X-100–insoluble fraction was higher in cells seeded with control fibrils compared with buffer-treated cells (Fig. 3A), while total TAU levels in the Triton X-100–soluble fraction remained unchanged across conditions (Fig. S9).

**Figure 3.**
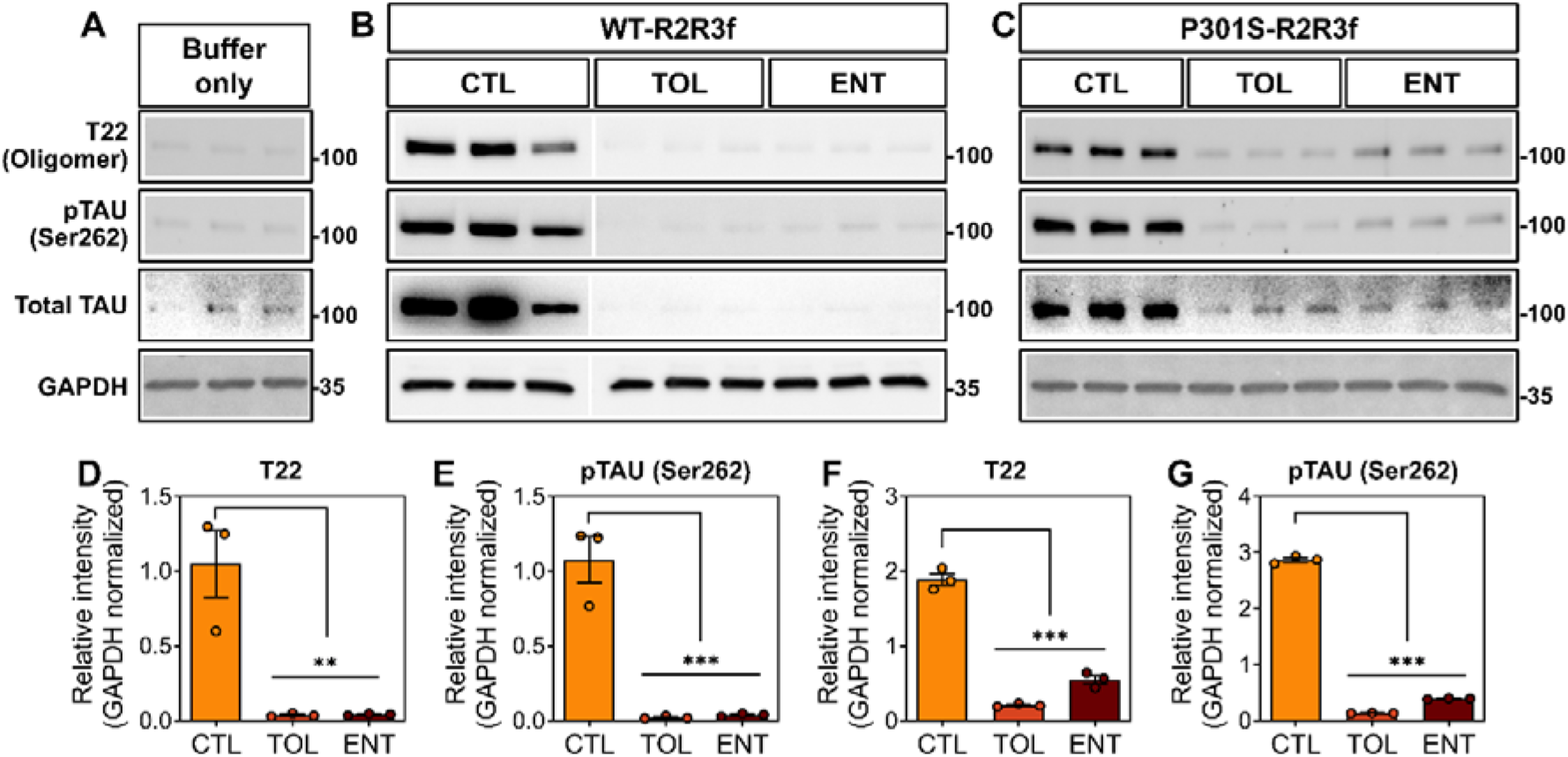
Effect of TOL- and ENT-modified R2R3f on TAU pathology in SY5Y-TAUP301L cells. (**A-C**) Western blot analysis of Triton X-100–insoluble fractions from differentiated SY5Y-TAUP301L cells treated with transfection buffer only (A) or transduced with WT-R2R3f (B) or P301S-R2R3f (C), formed in the absence of compounds (CTL) or in the presence of TOL or ENT (10 µM) during fibril formation. Representative blots of GAPDH from the soluble fraction are shown. Each lane represents an independent experiment (*n* = 3). Images of uncropped blots are provided in the Supplementary Information. (**D-G**) Quantification of TAU pathology in the Triton X-100–insoluble fraction. T22-positive oligomeric TAU (D, F) and pTAU (Ser262) (E, G) are shown for cells seeded with WT-R2R3f (D, E) or P301S-R2R3f (F, G) formed under CTL conditions or in the presence of TOL or ENT during fibril formation. Data are mean ± SEM (*n* = 3). \*\**P* < 0.01, \*\*\**P* < 0.001, one-way ANOVA with Dunnett’s multiple comparisons test.

Quantitative analysis confirmed significant reductions in T22-reactive oligomeric TAU and pTAU (Ser262) in cells seeded with WT-R2R3f or P301S-R2R3f formed in the presence of TOL or ENT compared with CTL fibrils (Fig. 3B–E). Unpaired statistical analyses directly comparing the relative effects of the compounds showed that, for P301S-R2R3f-seeded cells, TOL-modified fibrils were associated with significantly lower levels of T22-positive oligomeric TAU and pTAU (Ser262) compared with ENT-modified fibrils (Fig. S10). Differences between TOL and ENT were less pronounced for WT-R2R3f-seeded cells. Direct exposure of SH-SY5Y cells to TOL resulted in a concentration-dependent reduction in viability over 72 h, whereas ENT did not reduce viability below 50% at the highest concentration tested (Fig. S11).

### Sonicated K18 Fibrils are internalized by hiPSC-derived neurons and induce TAU aggregation and phosphorylation

To establish a TAU seeding paradigm in MAPT(P301L) isogenic human induced pluripotent stem cell-derived neurons (hiPSC-iNs), week-4 hiPSC(P301L)-iNs were exposed to sK18f labeled with Alexa Fluor 647 or to Alexa 647 dye alone (Fig. 4A). Week-4 hiPSC(P301L)-iNs were previously characterized as mature and phenotypically stable (Wu et al., 2019). Twenty-four hours after exposure, intracellular 647-positive puncta were detected within the neuronal soma in neurons treated with sK18f-647 but not in dye-only controls (Fig. 4B).

**Figure 4.**
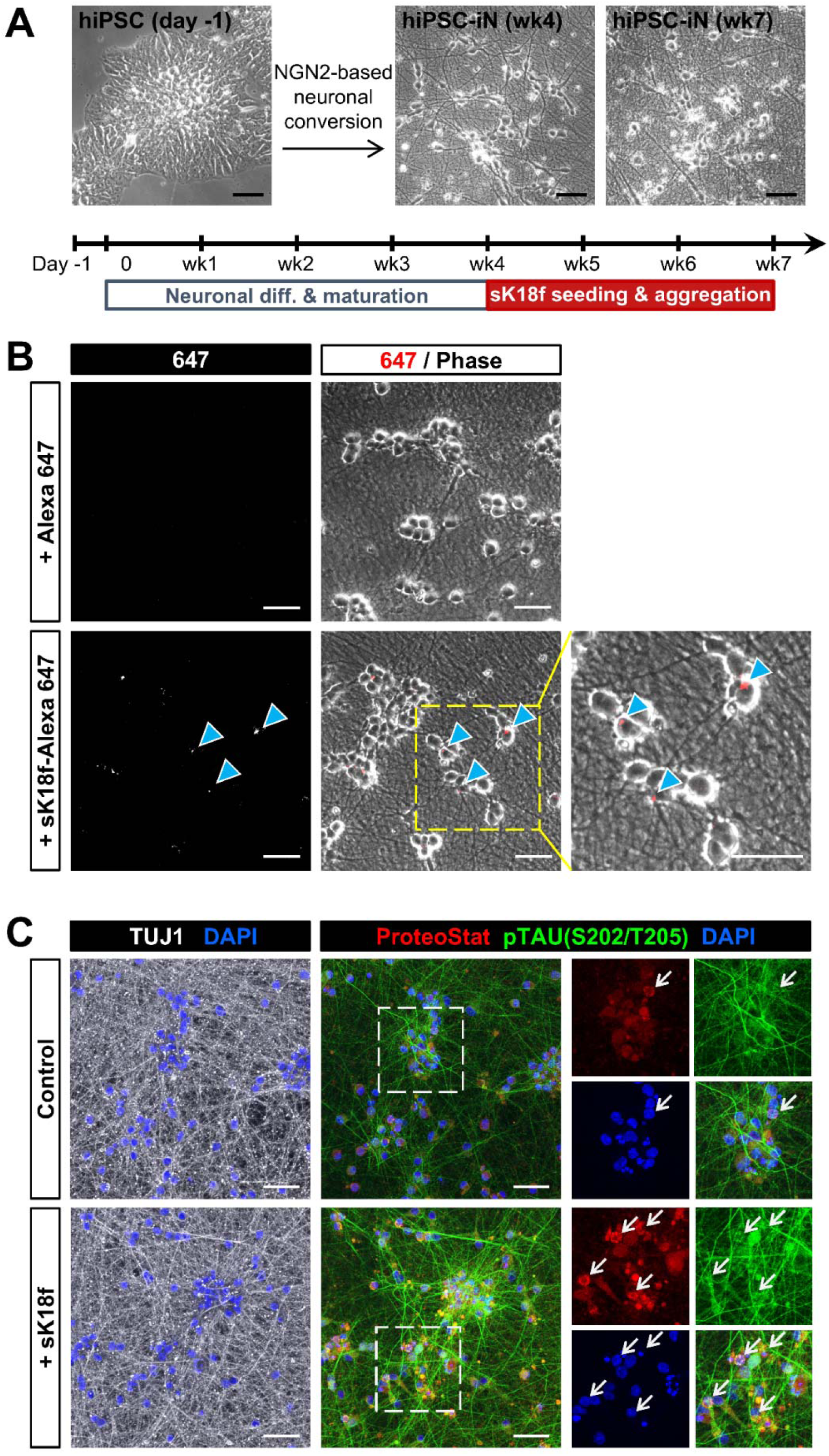
sK18 fibrils enter hiPSC-derived neurons and induce TAU aggregation. (**A**) Schematic of the experimental timeline showing NGN2-driven neuronal differentiation of hiPSCs and subsequent sK18f seeding phase (weeks 4–7). Representative phase-contrast images show the morphology of iPSCs, week-4 induced neurons, and week-7 mature neurons. Scale bar, 50 μm. (**B**) Alexa Fluor 647–labeled sK18 fibrils (sK18f-647) or Alexa Fluor 647 control dye were delivered into week-4 hiPSC-iNs using Xfect transfection. Images were acquired 24 h after transfection. The 647-fluorescence channel and merged 647/phase images are shown. Cyan arrowheads indicate 647-positive puncta within the dashed yellow box, with the corresponding high-magnification view displayed on the right. Scale bar, 50 μm. (**C**) hiPSC(P301L)-iNs were fixed at week 7 and immunostained for TUJ1, ProteoStat, pTAU(S202/T205), and DAPI. Full-field images of control and sK18f-treated cultures are shown. Dashed boxes indicate regions presented at higher magnification on the right, displaying individual channels for ProteoStat, pTAU(S202/T205), and DAPI. White arrows mark cells containing ProteoStat-positive puncta. Scale bar, 50 μm.

Following a three-week exposure to unlabeled sK18f, immunofluorescence analysis at week 7 showed increased ProteoStat-positive puncta and elevated pTAU(S202/T205) immunoreactivity in sK18f-treated cultures compared with control (Fig. 4C). ProteoStat-positive puncta were localized within the cytoplasm, and pTAU(S202/T205) signal was detected in TUJ1-positive neurons. Cultures showing increased ProteoStat puncta also exhibited overlapping pTAU(S202/T205) immunoreactivity.

### TOL and ENT modulate sarkosyl-insoluble TAU species in hiPSC-derived neurons

We next examined whether TOL or ENT modulates insoluble TAU species in neurons showing pre-existing TAU pathology. Sarkosyl fractionation followed by Western blot analysis was performed three weeks after sK18f exposure to assess TAU species in hiPSC(P301L)-derived neurons (Fig. 5A). sK18f treatment increased levels of pTAU(S202/T205), pTAU(S262), and total TAU in the sarkosyl-insoluble fraction, while corresponding changes in whole-cell lysates were minimal (Fig. 5B, C). LIVE/DEAD staining and LDH cytotoxicity assays showed no overt cytotoxicity in hiPSC-iNs treated with vehicle, TOL, or ENT (Fig. S12).

**Figure 5.**
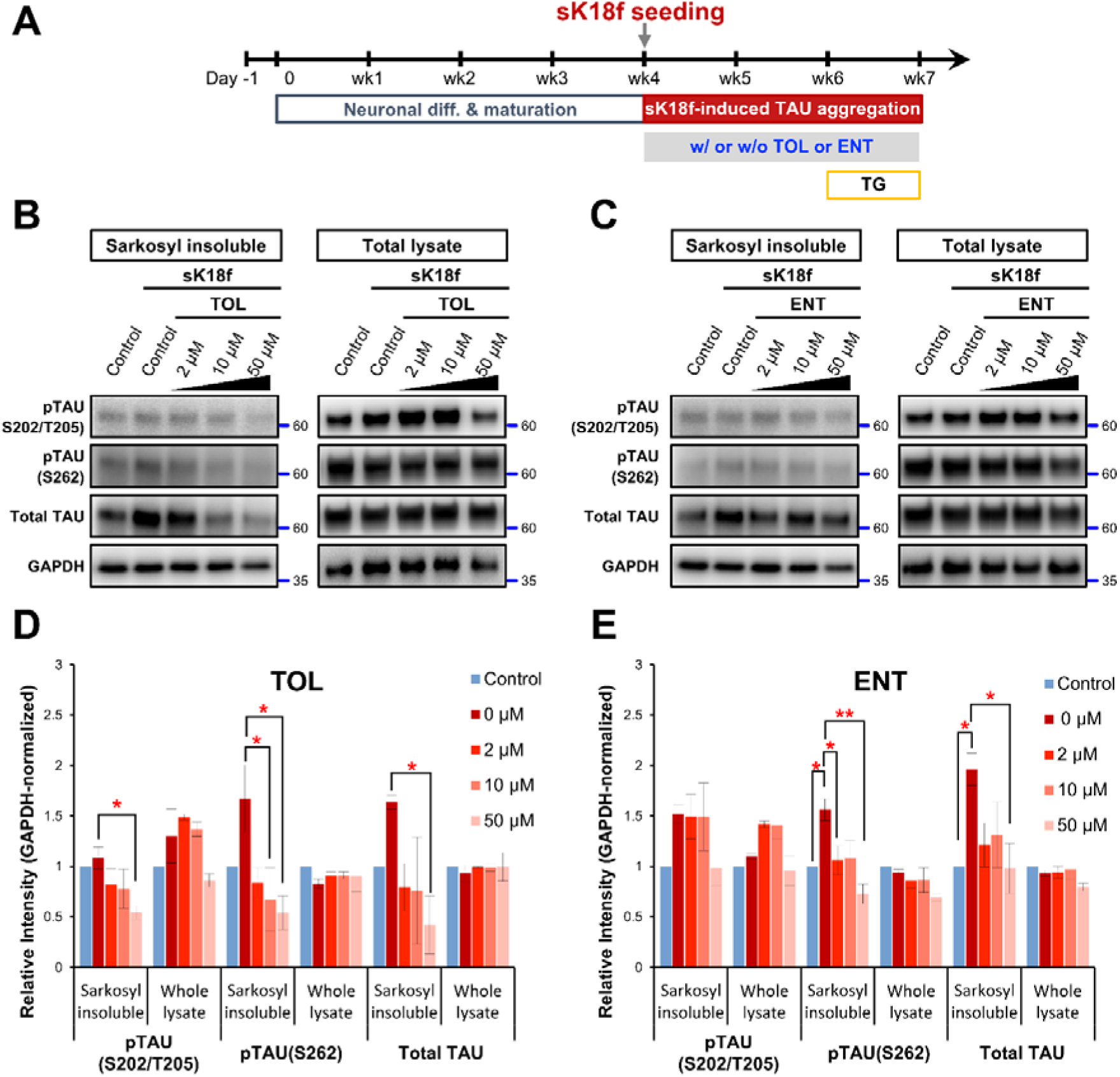
Effects of TOL and ENT on sarkosyl-insoluble TAU species in hiPSC(P301L)-derived neurons. (**A**) Experimental workflow showing the timeline of NGN2-induced neuronal differentiation, sK18f seeding, and treatment with TOL or ENT. Compounds were added one day after sK18f exposure and maintained for three weeks. Thapsigargin (TG) was applied for 1 week at week 6. (**B**, **C**) Western blots of pTAU(S202/T205), pTAU(S262), total TAU, and GAPDH in sarkosyl-insoluble fractions and total lysates from neurons treated with sK18f in the presence or absence of TOL (B) or ENT (C) at 2, 10, or 50 μM. Images of uncropped blots are provided in the Supplementary Information. (**D**, **E**) Quantification of GAPDH-normalized TAU signals in the sarkosyl-insoluble fraction and whole lysate corresponding to panels B and C. Data are mean ± SEM (*n* = 3 biological repeats), \**P* < 0.05, \*\**P* < 0.01, one-way ANOVA with Dunnett’s multiple comparisons test.

TOL or ENT was added 1 day after sK18f exposure and maintained throughout the incubation period, with thapsigargin applied during week 6 across all conditions as a uniform sensitizing background (Fig. 5A). TOL treatment reduced sarkosyl-insoluble pTAU and total TAU in a dose-dependent manner, with the largest reduction observed at the highest concentration tested (Fig. 5B, D). ENT treatment also reduced levels of insoluble TAU species, although the magnitude of change was consistently smaller across concentrations compared with TOL (Fig. 5C, E). Quantitative normalization of band intensities relative to unseeded vehicle controls confirmed that both TOL and ENT reduced pTAU(S262) and total TAU below the levels observed in the sK18f-seeded vehicle groups (Fig. 5D, E). In both treatment conditions, TAU levels in whole-cell lysates remained largely unchanged.

### TOL and ENT suppress sK18f-induced TAU aggregation and phosphorylation in hiPSC(P301L)-derived neurons

Immunofluorescence analysis was performed in hiPSC(P301L)-derived neurons following sK18f seeding at week 5 and subsequent compound treatment (Fig. 6A). sK18f treatment increased intracellular protein aggregation and TAU phosphorylation, indicated by elevated ProteoStat staining and increased pTAU (S202/T205) immunoreactivity compared with unseeded CTLs (Fig. 6B). Treatment with TOL or ENT reduced ProteoStat signal and pTAU (S202/T205) immunoreactivity relative to sK18f-treated neurons (Fig. 6B).

**Figure 6.**
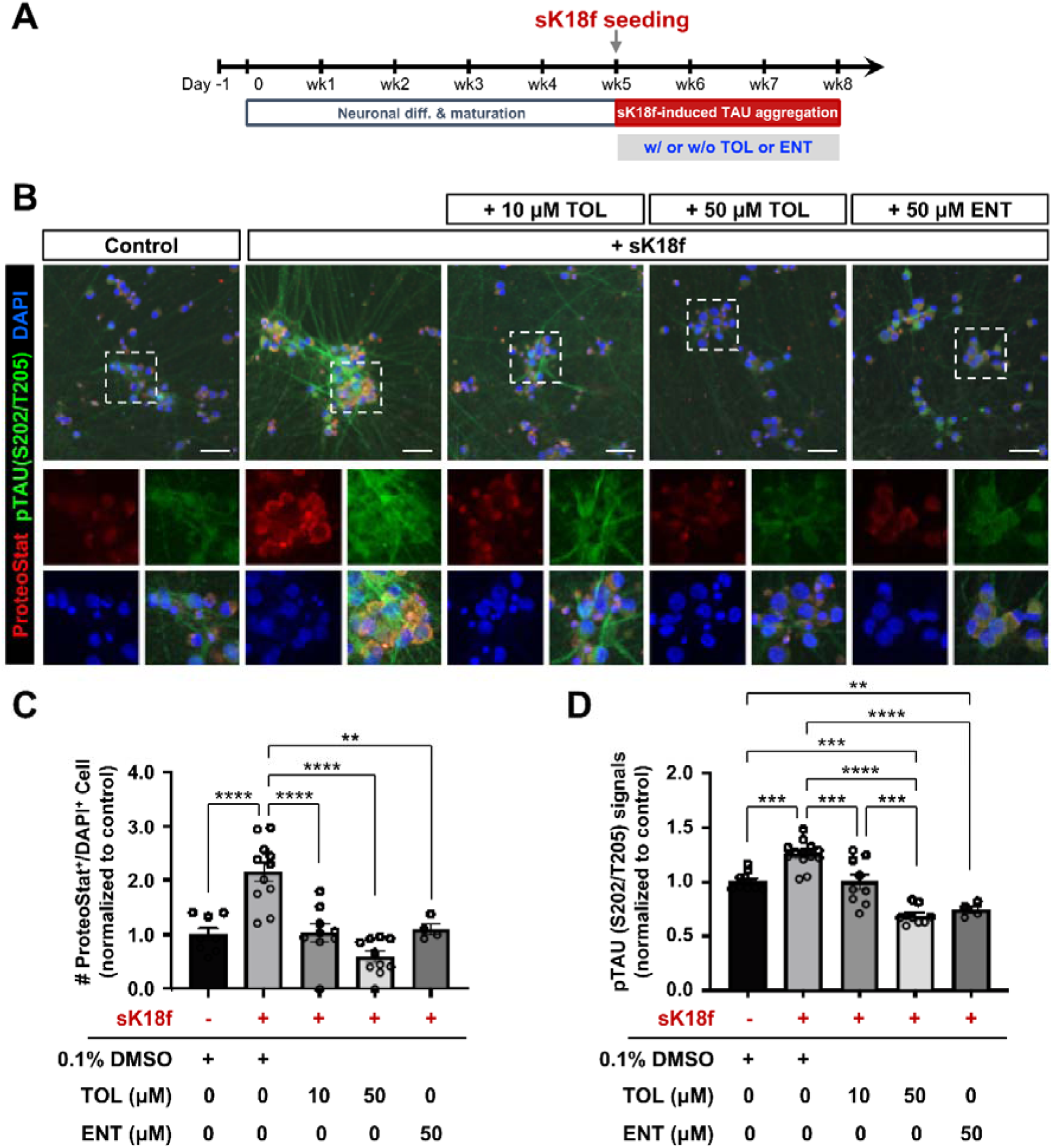
TOL and ENT reduce sK18f-induced TAU aggregation and phosphorylation in hiPSC(P301L)-derived neurons. **(A)** Timeline of sK18f seeding and drug treatment. TOL or ENT was added one day after sK18f exposure and maintained for three weeks. **(B)** Immunofluorescence of hiPSC-iNs fixed at week 8 under control conditions, sK18f treatment, or sK18f plus TOL (10 or 50 μM) or ENT (50 μM). Dashed boxes denote regions shown in magnified insets. Scale bar, 50 μm. **(C)** ProteoStat□/DAPI□cell counts normalized to control for TOL or ENT treatment conditions. Data are mean ± SEM (*n* = 4–12 imaging fields from 2 independent differentiation and treatment experiments), **P < 0.01, ****P < 0.0001, one-way ANOVA with Tukey’s HSD post-hoc test. **(D)** pTAU(S202/T205) fluorescence intensity normalized to control for TOL or ENT treatment conditions. Data are mean ± SEM (*n* = 4–12 imaging fields from 2 independent differentiation and treatment experiments), \*\**P* < 0.01, \*\*\**P* < 0.001, \*\*\*\**P* < 0.0001, one-way ANOVA with Tukey’s HSD post-hoc test.

Quantitative analysis showed a decrease in the number of ProteoStat^+^/DAPI^+^ cells and a reduction in pTAU (S202/T205) fluorescence intensity following compound treatment, normalized to unseeded controls (Fig. 6C, D). For TOL, a greater reduction in both readouts was observed at the higher concentration compared with the lower concentration (Fig. 6C, D). For TOL, a greater reduction in both readouts was observed at 50 µM compared with 10 µM (Fig. 6C). ENT treatment also decreased ProteoStat positivity and pTAU (S202/T205) immunoreactivity relative to sK18f-treated neurons, with a smaller magnitude of change than that observed with TOL (Fig. 6D).

## DISCUSSION

Tauopathies are driven by the accumulation and propagation of seed-competent TAU assemblies that template aggregation in recipient cells and promote the spread of pathology. Disease-associated *MAPT* mutations, particularly at residue P301, strongly enhance TAU aggregation and seeding, producing assemblies that differ structurally and functionally from WT TAU(Fischer et al., 2009; Chen et al., 2019). Identifying compounds that selectively target these pathogenic TAU conformers remains a key challenge in efforts to limit TAU-driven neurodegeneration.

In this study, we demonstrate that the clinically approved COMT inhibitors TOL and ENT suppress TAU aggregation in a concentration-dependent manner, exhibiting markedly greater potency against P301S mutant TAU than against WT TAU. Prior work has shown that catechol-containing compounds, including TOL and ENT, can inhibit the aggregation of TAU-derived peptides such as VQIVYK in cell-free systems, identifying the nitrocatechol moiety as a pharmacophore capable of interfering with amyloid formation (Mohamed et al., 2013). However, those studies did not assess whether aggregation inhibition differs between wild-type and disease-associated mutant TAU conformers. Our findings extend this earlier work by proving that catechol-based inhibition of TAU aggregation is strongly mutation-dependent. This aligns with structural evidence that P301 mutations destabilize the local architecture at the R2–R3 interface, thereby increasing the exposure of amyloidogenic motifs (Mirbaha et al., 2018; Chen et al., 2019).

Importantly, the suppression of bulk TAU aggregation does not necessarily translate into reduced TAU propagation. Therefore, it is critical to determine whether compounds that inhibit fibril formation also reduce the pathogenic activity of TAU assemblies in cellular systems. Our findings address a major limitation of many TAU aggregation inhibitors, which successfully reduce fibril formation in vitro but fail to block the seed-competent TAU species that actually drive the spread of pathology (Soeda et al., 2019; Soeda and Takashima, 2020; Hill et al., 2022).

SPR analysis showed that both TOL and ENT bind P301S TAU with higher affinity than wild-type TAU, providing a molecular correlation for their preferential functional effects on mutant TAU. Consistent with the SPR measurements, docking studies supported differential interactions of TOL and ENT with mutant versus wild-type TAU. Docking and molecular dynamics analyses indicated more favorable predicted binding for P301 mutant (P301S and P301L) TAU compared with WT, with TOL showing a modest advantage over ENT. These computational findings align with the stronger functional effects observed for TOL in cellular and neuronal models, supporting a structural basis for mutation-dependent compound activity without implying that binding affinity alone determines aggregation or seeding outcomes.

Indeed, SPR-derived affinities to monomeric TAU do not necessarily predict functional potency in aggregation or seeding assays (Schafer et al., 2013; Chai et al., 2024). This is because aggregation can also be modulated through transient interactions with intermediate species or the redirection of aggregation pathways. High-resolution cryo-electron microscopy studies have shown that pathogenic TAU fibrils share a conserved core structure centered on the VQIVYK motif (Fitzpatrick et al., 2017; Falcon et al., 2018). Therefore, the enhanced binding of catechol-containing compounds to the P301 mutant TAU may reflect increased accessibility of interaction surfaces within these aggregation-prone regions, even though seeding inhibition remains a dynamic, multistep process that extends beyond simple binding affinity.

The functional relevance of these findings was further supported in neuronal models. In SY5Y-TAUP301L cells, fibrils formed in the presence of TOL or ENT induced lower levels of TAU oligomerization and Ser262 phosphorylation, indicating reduced pathogenic activity of the resulting assemblies. By testing the compounds during fibril formation, this experiment demonstrates that drug-modified TAU assemblies exhibit reduced pathogenic capacity upon entering recipient cells. Catechol-based compounds have previously been reported to reduce TAU oligomerization and pathological TAU accumulation in cellular systems and transgenic mouse models through cysteine-dependent mechanisms (Soeda et al., 2015). However, those studies focused primarily on early oligomer formation and overall pathology burden, rather than directly examining seed competence or post-seeding propagation. Our results extend this work by demonstrating that catechol-containing compounds can reduce the downstream pathological impact of TAU assemblies with defined seeding histories.

Extending these observations to a human neuronal context, we leveraged human iPSC-derived neuronal seeding models to test translational relevance. We showed that pathogenic K18 fibrils are efficiently internalized by hiPSC-derived neurons, inducing robust TAU aggregation and phosphorylation that recapitulate key features of tauopathy (Goedert and Spillantini, 2017). In this system, TOL and ENT were applied after fibril exposure to determine whether these compounds can suppress intracellular TAU pathology once seeding has been initiated. Treatment with TOL, and to a lesser extent ENT, reduced sarkosyl-insoluble TAU species and cellular markers of aggregation and phosphorylation. Notably, we incorporated orthogonal viability controls (LIVE/DEAD staining and LDH release) under vehicle-matched conditions, supporting that the observed reductions in seeded TAU readouts are not readily explained by overt cytotoxicity. The consistency of compound effects across biochemical assays, cellular models, and human iPSC-derived neurons, particularly in P301 mutant contexts, supports a model in which TOL and ENT preferentially interfere with mutation-driven, seed-competent TAU assemblies.

From a translational perspective, the clinical pharmacology of TOL and ENT is directly relevant to their observed effects. TOL crosses the blood–brain barrier and inhibits COMT centrally, whereas ENT acts primarily as a peripheral COMT inhibitor with limited brain penetration (Lees, 2008). This distinction is consistent with the stronger neuronal effects observed for TOL in the present study. At the same time, the known hepatotoxicity associated with TOL limits its direct repurposing for chronic use (Artusi et al., 2021). Instead, these findings highlight catechol-based scaffolds as starting points for the development of TAU-directed compounds with improved safety and central exposure.

More broadly, our results underscore the importance of mutation-specific strategies targeting seed-competent TAU species, which are key drivers of pathological spread in tauopathies. This study is not without limitations. First, our human iPSC-neuronal validation was performed using a single MAPT(P301L) isogenic clone; future work must replicate these findings across independently edited clones or patient-derived lines to ensure generalizability. Second, while our data clearly show endpoint reduction in pathology, further studies are required to define the precise structural mechanisms by which these compounds interact with transient aggregation intermediates. Finally, while our human iPSC models are highly relevant to human disease, future studies must test optimized versions of these compounds in more complex systems, such as 3D brain organoids or animal models.

In conclusion, our findings show that clinically approved COMT inhibitors, TOL and ENT, preferentially inhibit the aggregation of pathogenic P301 mutant TAU and reduce the downstream pathogenic activity of TAU assemblies in neuronal systems. By demonstrating mutation-dependent suppression of aggregation and seeded intracellular TAU pathology across biochemical assays, cellular models, and human iPSC-derived neurons, this study validates the pharmacological modulation of seed-competent TAU assemblies as a viable therapeutic strategy. These results directly support the development of catechol-based scaffolds targeting pathogenic TAU conformers to halt tauopathy progression.

## DECLARATIONS

### Ethics Approval

Not applicable

### Data Availability Statement

All data supporting the findings of this study are provided within the article and its Supplementary Information. In addition, all raw and processed datasets are publicly available at Zenodo (DOI: https://doi.org/10.5281/zenodo.18618834). The repository includes raw ThT kinetic traces, TEM fibril width measurements, ImageJ macros, global kinetic-fitting analyses, and complete MM-GBSA and per-residue energy-decomposition outputs.

### Consent to Participate

Not applicable.

### Consent for Publication

Not applicable.

### Author Contributions

**Methodology**: Ihor Kozlov, Yu-Sheng Hung; **Formal analysis and investigation**: Ihor Kozlov, Yu-Sheng Hung, Alladi Charanraj Goud, Roman Kouřil, Sudeep Roy, Yu-Hui Wong, Viswanath Das; **Writing – original draft preparation**: Ihor Kozlov, Yu-Sheng Hung, Sudeep Roy, Yu-Hui Wong, Viswanath Das; **Writing – review and editing**: Yu-Hui Wong, Viswanath Das; **Funding acquisition**: Yu-Hui Wong, Viswanath Das; **Resources**: Yu-Hui Wong, Viswanath Das; **Supervision**: Yu-Hui Wong, Viswanath Das; **Conceptualization**: Viswanath Das.

## Acknowledgements

The research was funded by the Grant Agency of the Czech Republic (GAČR, 23-06301J). Support was also provided by the National Institute for Neurological Research (Program EXCELES, ID Project No. LX22NPO5107), funded by the European Union—Next Generation EU through the Ministry of Education, Youth, and Sports of the Czech Republic (MEYS). Additional contributions were supported by the infrastructural projects CZ-OPENSCREEN (LM2023052) and EATRIS-CZ (LM2023053), the Czech biobank network BBMRI (LM2023033) for biobanking resources, and the Czech-BioImaging (LM2023050, LM2018129) funded by MEYS, and the project TN02000109 (Personalized Medicine: From Translational Research into Biomedical Applications) by the Technology Agency of the Czech Republic. IH acknowledges the Internal Student Grant Agency of Palacký University in Olomouc, Czech Republic (IGA_LF_2024_038).

