## Supplementary Material for "Repurposed COMT Inhibitors Tolcapone and Entacapone Selectively Suppress Aggregation and Seeding of P301 Mutant TAU in Human Neuronal Models"

### **Table of Content**

**Section A. Supplementary Figures**

- **Figure S1**. Concentration-dependent suppression of tau aggregation kinetics by tolcapone (TOL) and entacapone (ENT).
- **Figure S2**. Multi-Injection Cycle Kinetic (MICK) surface plasmon resonance (SPR) sensorgrams showing binding of TOL and ENT to WT and P301S R2R3 tau constructs.
- **Figure S3**. AlphaFold2-predicted structures of WT, P301L, and P301S tau colored by per‑residue confidence.
- **Figure S4**. Per‑residue AlphaFold2 pLDDT confidence scores for WT, P301L, and P301S tau.
- **Figure S5**. AlphaFold2 predicted aligned error (PAE) maps for WT, P301L, and P301S tau.
- **Figure S6**. AlphaFold2 MSA sequence coverage profiles for WT, P301L, and P301S tau.
- **Figure S7**. Ramachandran plots of backbone dihedral angle distributions for WT, P301L, and P301S tau models.
- **Figure S8**. Docking interactions of entacapone (ENT) and tolcapone (TOL) with the P301L tau filament model.
- **Figure S9**. Soluble tau levels are not altered following seeding with compound-modified R2R3 fibrils.
- **Figure S10**. Comparison of tau pathology induced by tolcapone (TOL) or entacapone (ENT)-modified R2R3 fibrils.
- **Figure S11**. Tolcapone (TOL) or entacapone (ENT) does not cause overt cytotoxicity within the assay window.
- **Figure S12**. TOL and ENT do not induce overt cytotoxicity in hiPSC-derived neurons

**Section B. Supplementary Tables**

- **Table S1**. Induced-fit docking scores for ENT and TOL against WT, P301L, and P301S tau.
- **Table S2**. MM-GBSA binding free energy decomposition for ENT–WT.
- **Table S3**. MM-GBSA binding free energy decomposition for TOL–WT.
- **Table S4**. MM-GBSA binding free energy decomposition for ENT–P301S.
- **Table S5**. MM-GBSA binding free energy decomposition for TOL–P301S
- **Table S6**. MM-GBSA binding free energy decomposition for ENT–P301L.
- **Table S7**. MM-GBSA binding free energy decomposition for TOL–P301L.

**Section C. Supplementary Methods**

- SH-SY5Y Cell Viability Assay
- Homology Modeling
- Molecular docking
- Molecular dynamics simulations and binding free energy calculations
- Per-Residue Decomposition
- Supplementary References

**Section D. Uncropped Western Blots**

- Uncropped images corresponding to the main Figures

### **Section A (Supplementary Figures)**

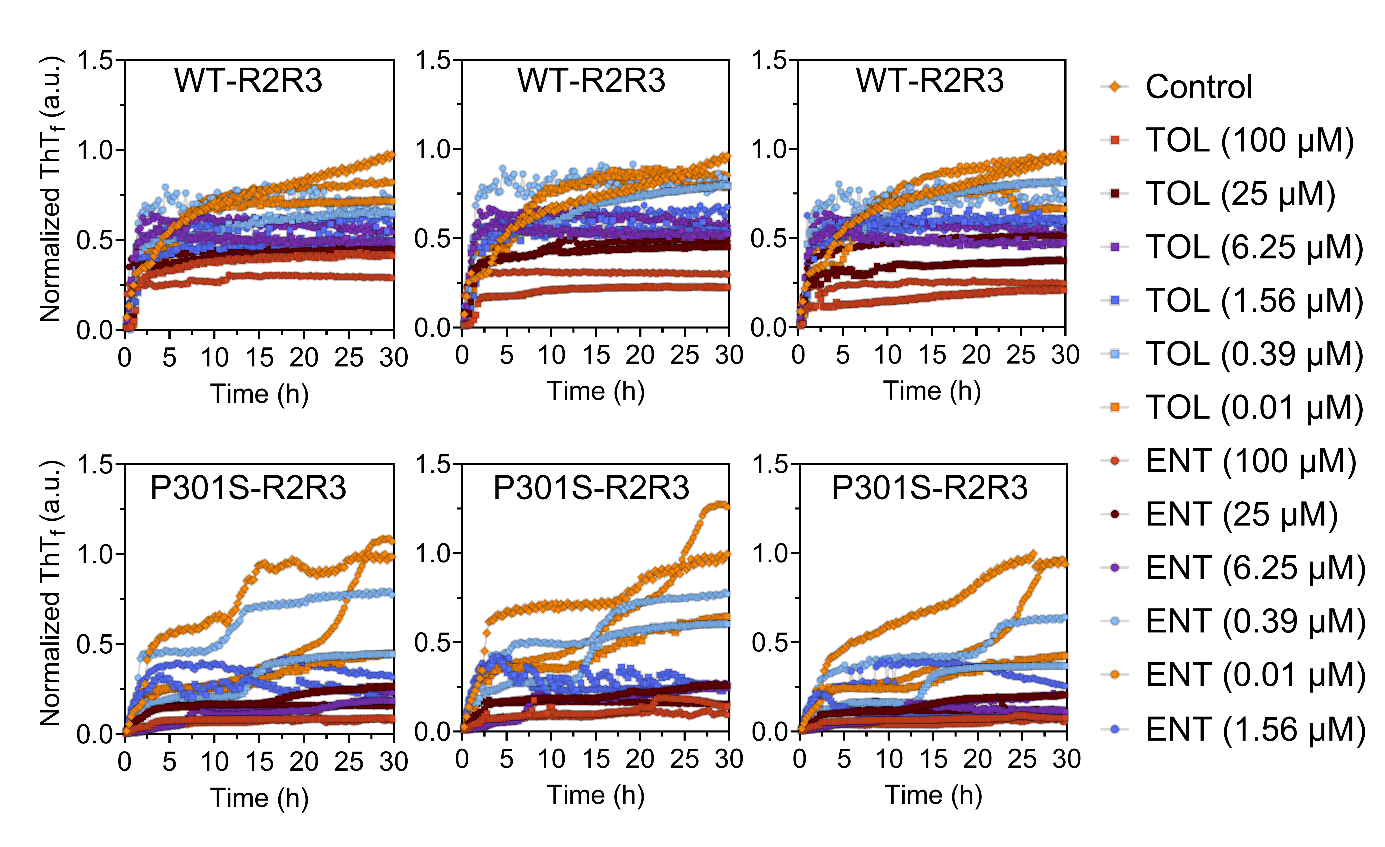
**Figure S1.** **Concentration-dependent suppression of tau aggregation kinetics by tolcapone (TOL) and entacapone (ENT).**

Thioflavin T (ThT) fluorescence kinetic traces showing the effects of increasing concentrations of tolcapone (TOL) and entacapone (ENT) on the aggregation of WT-R2R3 and P301S-R2R3 tau constructs.

Under control conditions, both constructs exhibit characteristic sigmoidal aggregation kinetics, with P301S-R2R3 showing a faster onset and a higher ThT signal than WT-R2R3. Increasing concentrations of TOL or ENT progressively delay the onset of aggregation and reduce overall ThT fluorescence for both constructs, with more pronounced suppression observed for P301S-R2R3 at lower compound concentrations.

Aggregation reactions were performed in the presence of TOL or ENT at 100, 25, 6.25, 1.56, 0.39, or 0.01 µM, alongside untreated controls. The area under the curve (AUC) derived from these kinetic traces was used to quantify aggregation and to generate the concentration–response plots shown in Fig. 1G and Fig. 1H.

Data are shown as mean traces from three independent experiments. Raw ThT fluorescence values are available via Zenodo (Data Availability Statement).

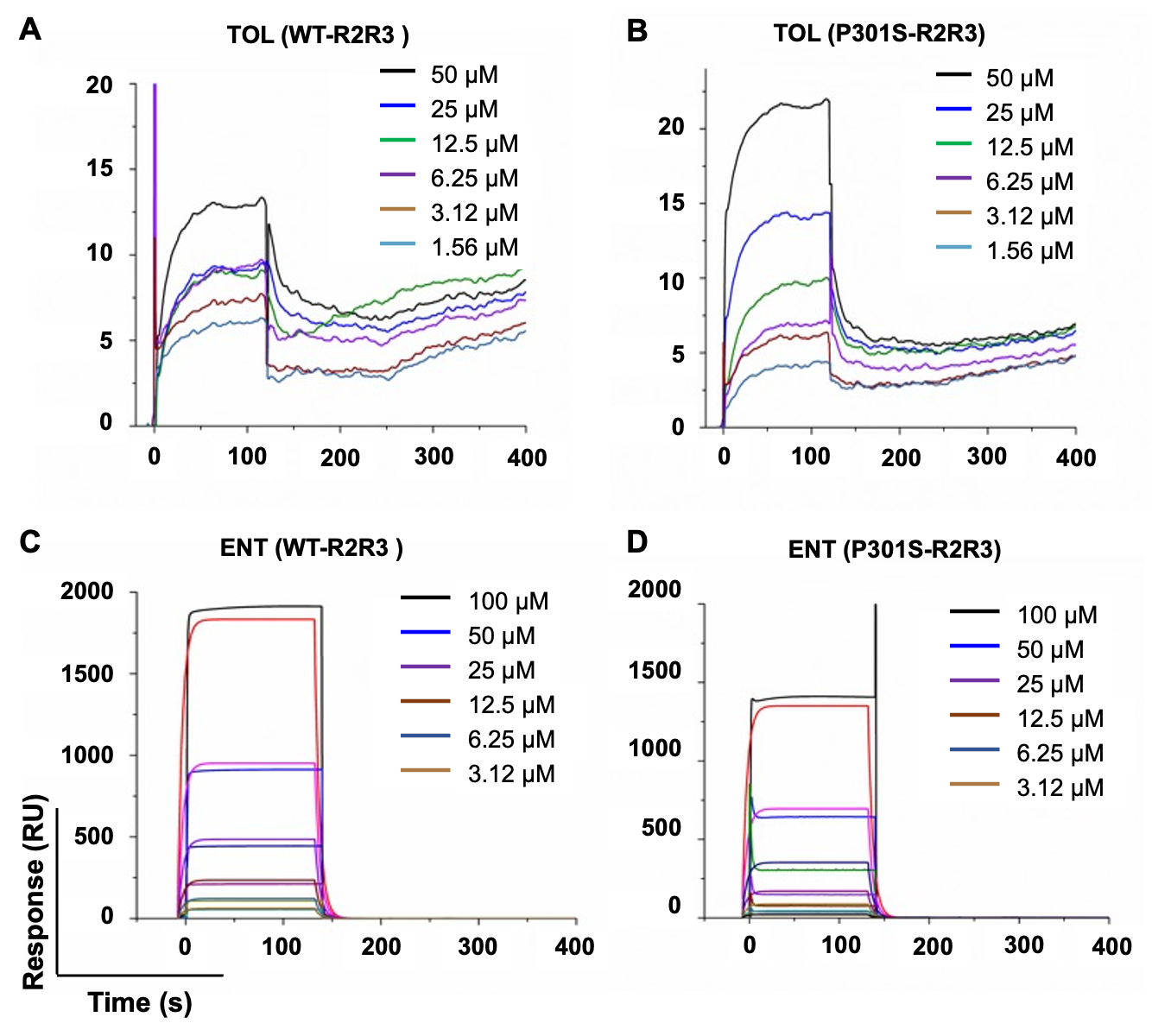

**Figure S2**. **Multi-Injection Cycle Kinetic (MICK) surface plasmon resonance (SPR) sensorgrams showing binding of TOL and ENT to WT and P301S R2R3 tau constructs.**

Surface plasmon resonance (SPR) sensorgrams showing the interactions of tolcapone (TOL) and entacapone (ENT) with monomeric WT-R2R3 and P301S-R2R3 tau constructs at a range of analyte concentrations.

(**A**, **B**) Binding responses of TOL to WT-R2R3 (A) and P301S-R2R3 (B) tau.

(**C**, **D**) Binding responses of ENT to WT-R2R3 (C) and P301S-R2R3 (D) tau.

R2R3 tau peptides were immobilized on the sensor surface at a constant density, and binding measurements were performed using a Multi Injection Cycle Kinetic (MICK) format with sequential injections of increasing concentrations of TOL or ENT. Each injection produced distinct association and dissociation phases, with higher analyte concentrations yielding proportionally larger response signals. Sensorgrams demonstrate reproducible, concentration-dependent binding for both compounds to WT and P301S tau constructs, with stronger responses observed for TOL and for P301S-R2R3 relative to WT-R2R3.

Kinetic and equilibrium binding parameters were obtained by global kinetic fitting of the full concentration series using a 1:1 interaction model and are summarized in Table 1. Source SPR data are available via Zenodo (see Data Availability Statement).

**Figure S3. AlphaFold2-predicted structures of WT, P301L, and P301S tau colored by per‑residue confidence.**

**(A)** Wild-type tau (WT), **(B)** P301L, and **(C)** P301S models predicted using AlphaFold2 are shown in cartoon representation and colored according to the per‑residue confidence score (pLDDT; 0–100), from blue (very high confidence, pLDDT > 90) through cyan (high confidence, 70–90), yellow (low confidence, 50–70) to orange/red (very low confidence, < 50). All three variants display a similar overall architecture within the structured core, whereas the N‑ and C‑terminal regions and flexible linkers are characterized by uniformly low pLDDT values, consistent with intrinsic disorder. Substitution of Pro301 with Leu or Ser does not markedly alter the global fold but induces subtle local rearrangements around residue 301 within regions predicted with intermediate–high confidence, suggesting that these mutations primarily modulate local packing and dynamics rather than causing large‑scale refolding of tau.

**Figure S4. Per‑residue AlphaFold2 pLDDT confidence scores for WT, P301L, and P301S tau.**

(**A**) Per‑residue pLDDT scores for WT tau**, (B)** P301L tau, and **(C)** P301S tau predicted by AlphaFold2. For each construct, the pLDDT score (0–100) is plotted against residue index, with horizontal reference bands indicating very high confidence (pLDDT > 90), high confidence (70–90), low confidence (50–70), and very low confidence (< 50). All three profiles show a conserved pattern: a central region with predominantly high pLDDT values, reflecting a well‑defined structural core, and extended N‑ and C‑terminal segments with low pLDDT values, indicative of intrinsic structural disorder. The similarity of the pLDDT traces between WT, P301L, and P301S suggests that the single‑point mutations at position 301 do not substantially alter the global confidence landscape of the models but instead are expected to induce only localized changes in structure and dynamics.

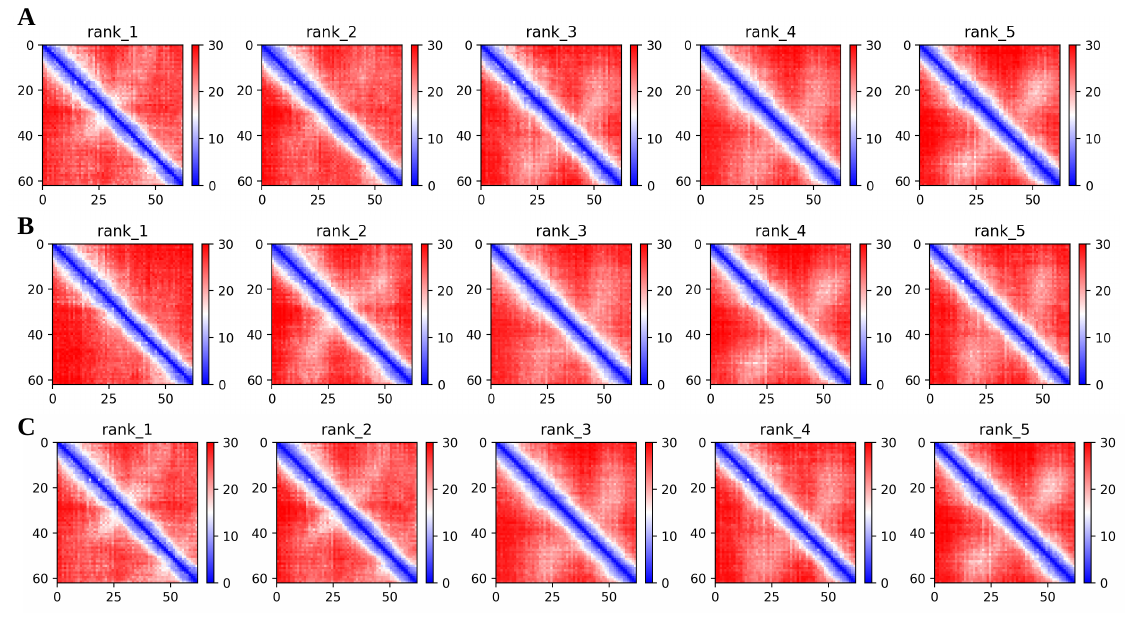

**Figure S5. AlphaFold2 predicted aligned error (PAE) maps for WT, P301L, and P301S tau.**

**(A)** WT tau, **(B)** P301L, and **(C)** P301S PAE maps calculated by AlphaFold2 are shown as 2D matrices, in which each pixel represents the predicted positional error (in Å) for residue i when the model is aligned on residue j. The color scale ranges from dark green (low error, < 5 Å; high confidence in the relative positioning of residue pairs) to yellow–red (high error, > 15–20 Å; low confidence in relative positioning). All three variants display relatively low PAE values along the main diagonal blocks, indicating reliable local geometry within contiguous sequence regions, and higher PAE values for many off‑diagonal blocks, reflecting uncertainty in the relative orientation of distant segments and supporting substantial inter‑domain and long‑range flexibility. The overall similarity of the PAE patterns for WT, P301L and P301S indicates that the single‑residue substitutions at position 301 do not markedly alter the predicted inter‑residue geometry at the global level, but instead are expected to affect local packing and dynamics in the vicinity of residue 301.

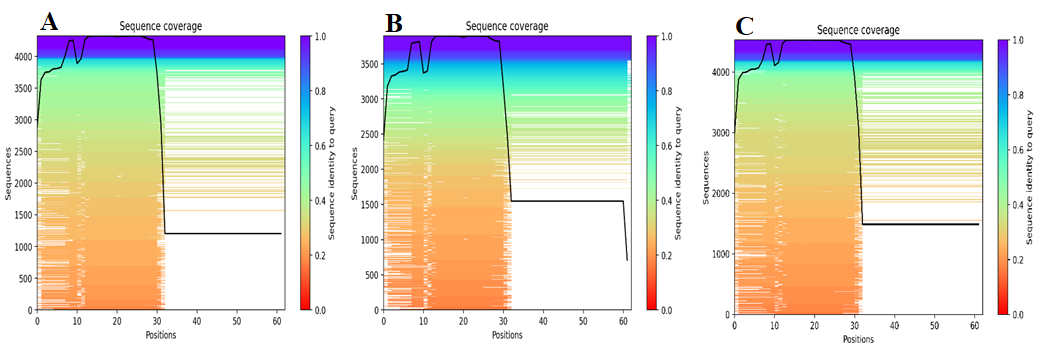

**Figure S6. AlphaFold2 MSA sequence coverage profiles for WT, P301L, and P301S tau.**

**(A)** WT tau, **(B)** P301L, and **(C)** P301S sequence coverage plots generated by AlphaFold2. For each construct, the number of aligned homologous sequences in the multiple sequence alignment (sequence coverage) is shown as a function of residue index. Regions with high coverage reflect positions that are well represented across homologs in the MSA, whereas dips in coverage highlight segments with limited evolutionary information. The three variants display nearly overlapping coverage patterns across the entire sequence, including the region around residue 301, demonstrating that the P301L and P301S substitutions do not alter the underlying MSA composition or depth used for structure prediction. This comparable sequence coverage supports a direct comparison of the AlphaFold2 confidence metrics (pLDDT and PAE) among WT, P301L, and P301S, as all three models are constrained by a similar level of evolutionary information.

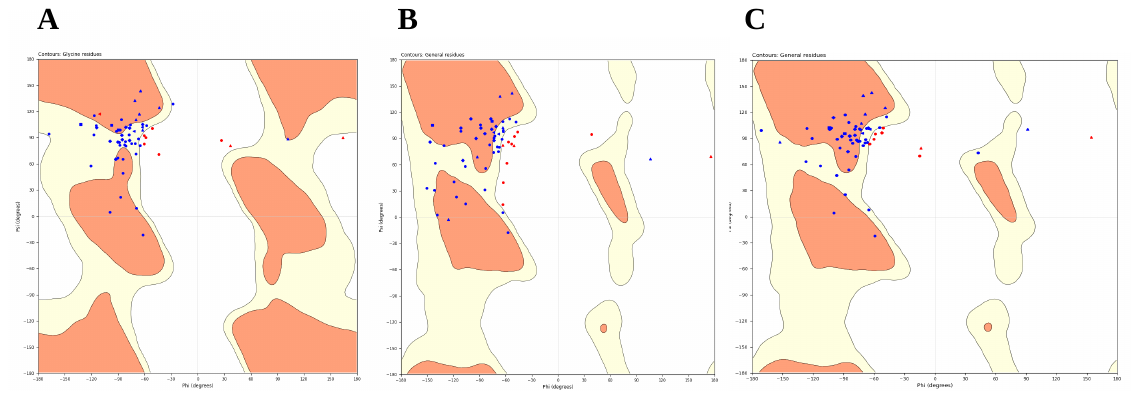

**Figure S7. Ramachandran plots of backbone dihedral angle distributions for WT, P301L, and P301S tau models.**

**(A)** WT tau, **(B)** P301L, and **(C)** P301S Ramachandran plots derived from the AlphaFold2-predicted structures. Each point represents the backbone ϕ and ψ angles of a non-glycine, non-proline residue, mapped onto the classical Ramachandran space with most favored, additionally allowed, and disallowed regions indicated. The majority of residues in all three models cluster within favored and allowed regions, with only a small number of residues falling into disallowed regions, consistent with good backbone stereochemistry. The similar distributions observed for WT, P301L, and P301S indicate that neither mutation produces substantial conformational outliers or unusual backbone strain at the global level, supporting the structural plausibility of the AlphaFold2 models used for subsequent analyses.

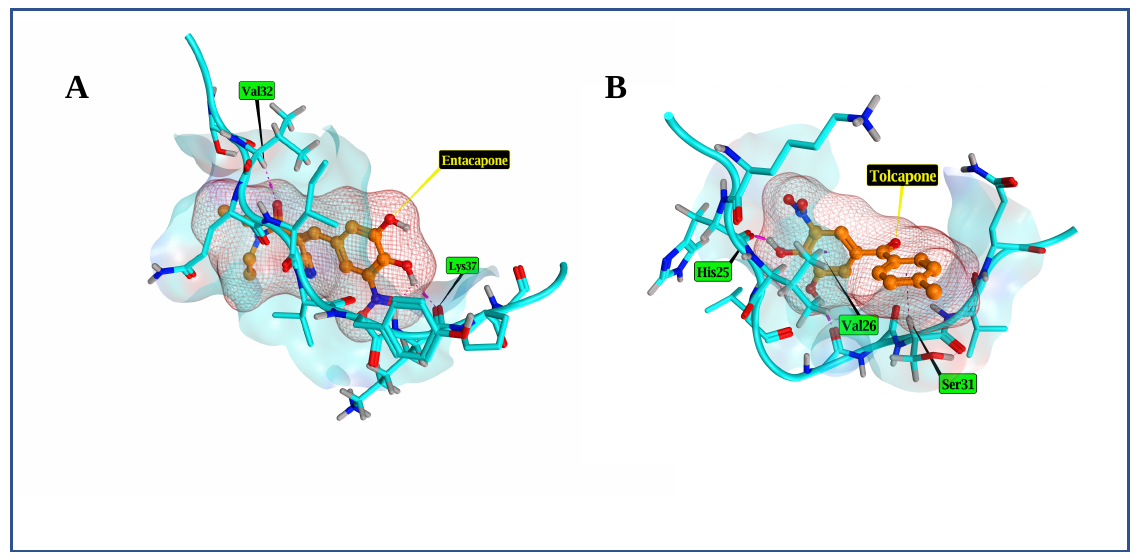

**Figure S8.** **Docking interactions of entacapone (ENT) and tolcapone (TOL) with the P301L tau filament model.**

Predicted binding poses of **(A)** entacapone (ENT) and **(B)** tolcapone (TOL) docked to the AlphaFold2‑derived P301L tau filament model. The tau filament is shown as a ribbon in surface representation, with key interacting residues highlighted as sticks. Ligands are depicted as space‑filling models colored by atom type. Docking was performed using MOE 2024.01, and interactions are indicated as follows: hydrogen bonds (dashed yellow lines), hydrophobic contacts (pink), and potential ionic interactions (violet lines). Both ligands occupy a common binding pocket on the filament surface, engaging in hydrogen bonds with polar side chains and π‑stacking or hydrophobic contacts with aromatic residues. The similar interaction patterns for ENT and TOL provide a structural basis for their comparable binding affinities to the P301L mutant as quantified by MM‑GBSA.

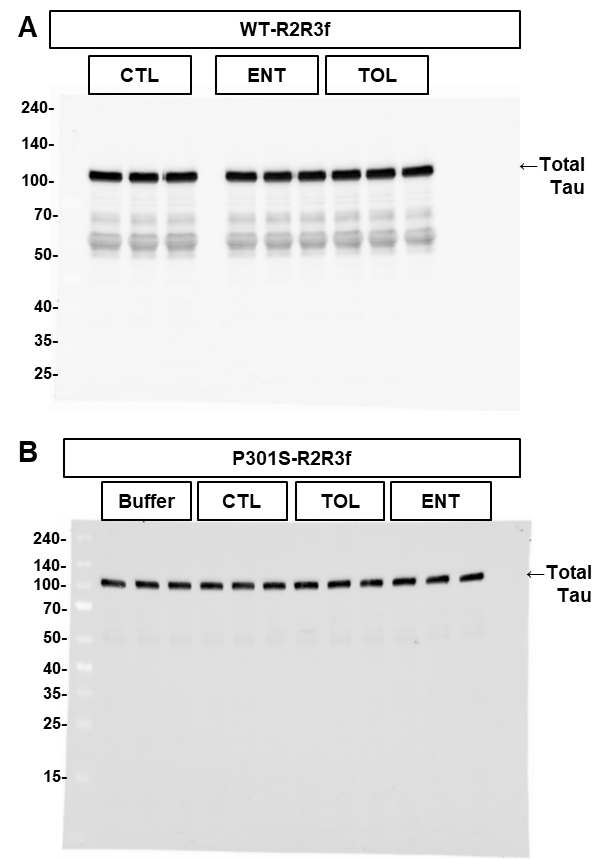

**Figure S9. Soluble tau levels are not altered following seeding with compound-modified R2R3 fibrils.**

Western blot analysis of Triton X-100–soluble fractions from differentiated SY5Y-TauP301L-EGFP cells transduced with WT-R2R3 fibrils (WT-R2R3f; A) or P301S-R2R3 fibrils (P301S-R2R3f; B) generated in the absence of compounds (CTL) or in the presence of tolcapone (TOL; 10 µM) or entacapone (ENT; 10 µM), or treated with transfection buffer only (buffer).

Across all conditions, levels of total tau in the Triton X-100–soluble fraction remain comparable, indicating that seeding with control or compound-modified fibrils does not measurably affect soluble tau abundance.

These data support the interpretation that changes observed in the Triton X-100–insoluble fraction reflect differences in aggregation and recruitment of tau into insoluble assemblies rather than alterations in overall tau expression. Each lane represents an independent experiment (*n* = 3).

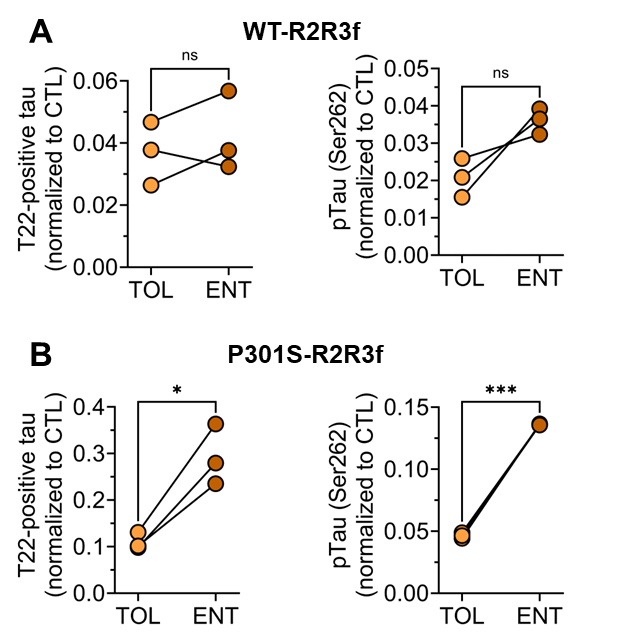

**Figure S10**. **Comparison of tau pathology induced by tolcapone (TOL) or entacapone (ENT)-modified R2R3 fibrils.**

Paired analysis of tau pathology in SY5Y-TauP301L-EGFP cells seeded with R2R3 fibrils formed in the presence of TOL or ENT.

**(A)** Cells seeded with WT-R2R3f, showing GAPDH-normalized levels of T22-positive oligomeric tau (left) and pTau (Ser262) (right).

**(B)** Cells seeded with P301S-R2R3f, showing GAPDH-normalized levels of T22-positive oligomeric tau (left) and pTau (Ser262) (right).

Each data point represents an independent biological replicate (*n* = 3). Signals are normalized to the corresponding CTL condition. *P < 0.05, ***P < 0.001, unpaired, two-tailed Student’s *t*-test.

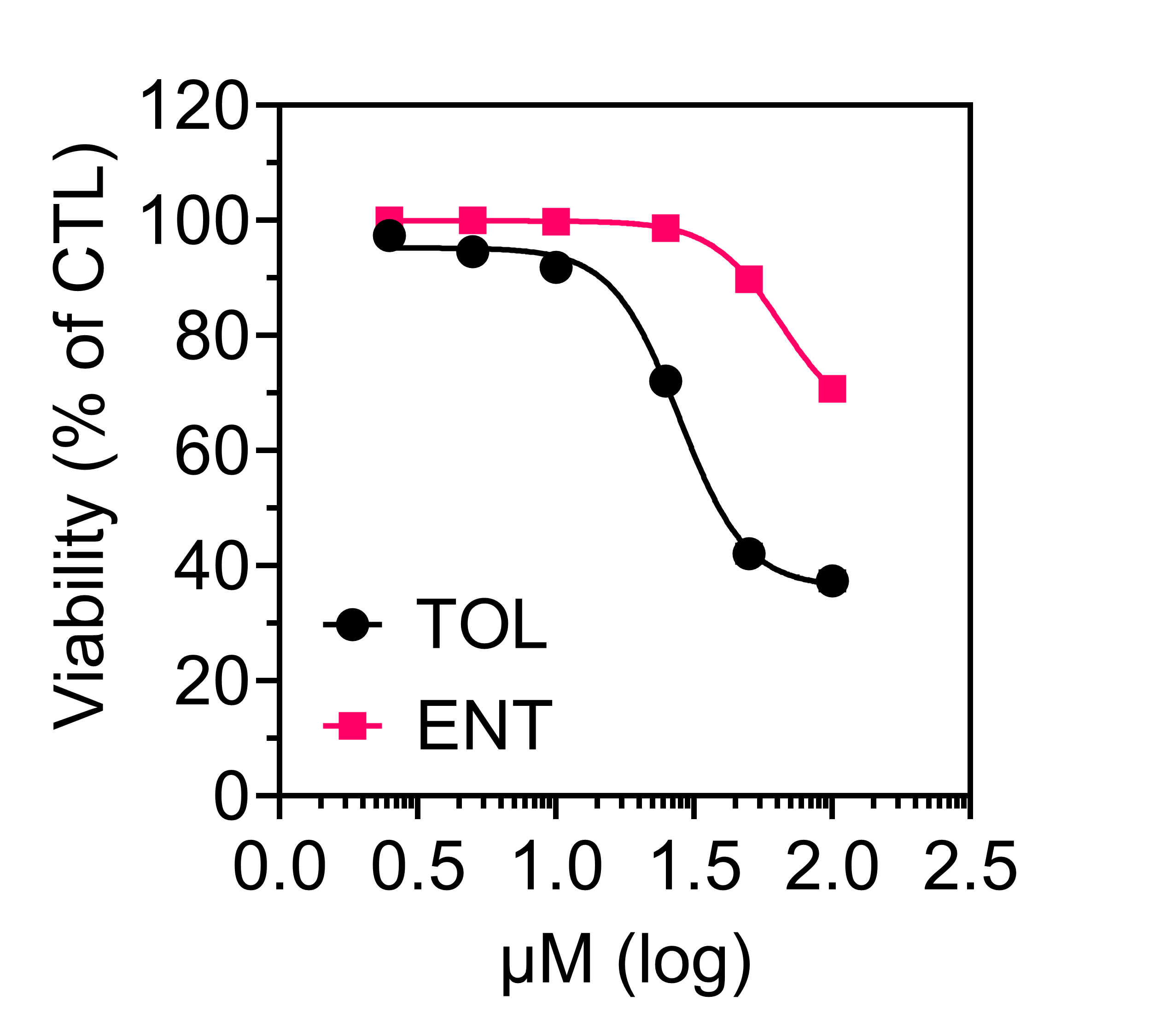

**Figure S11. Tolcapone (TOL) or entacapone (ENT) does not cause overt cytotoxicity within the assay window**.

Cell viability was assessed by MTS assay following exposure to TOL or ENT under conditions relevant to the functional assays used in this study.

SH-SY5Y cells were treated for 72 h. TOL reduced viability in a concentration-dependent manner and crossed the 50% viability threshold within the tested concentration range, allowing estimation of an IC_50_ by four-parameter logistic fitting. In contrast, ENT showed minimal effects on SH-SY5Y cell viability and did not reduce viability below 50% at the highest concentration tested; therefore, an IC_50_ was not determined under these conditions.

All data are normalized to vehicle controls and shown as mean ± SEM (*n* = 3). Raw absorbance values and nonlinear regression outputs are available via Zenodo (Data Availability Statement).

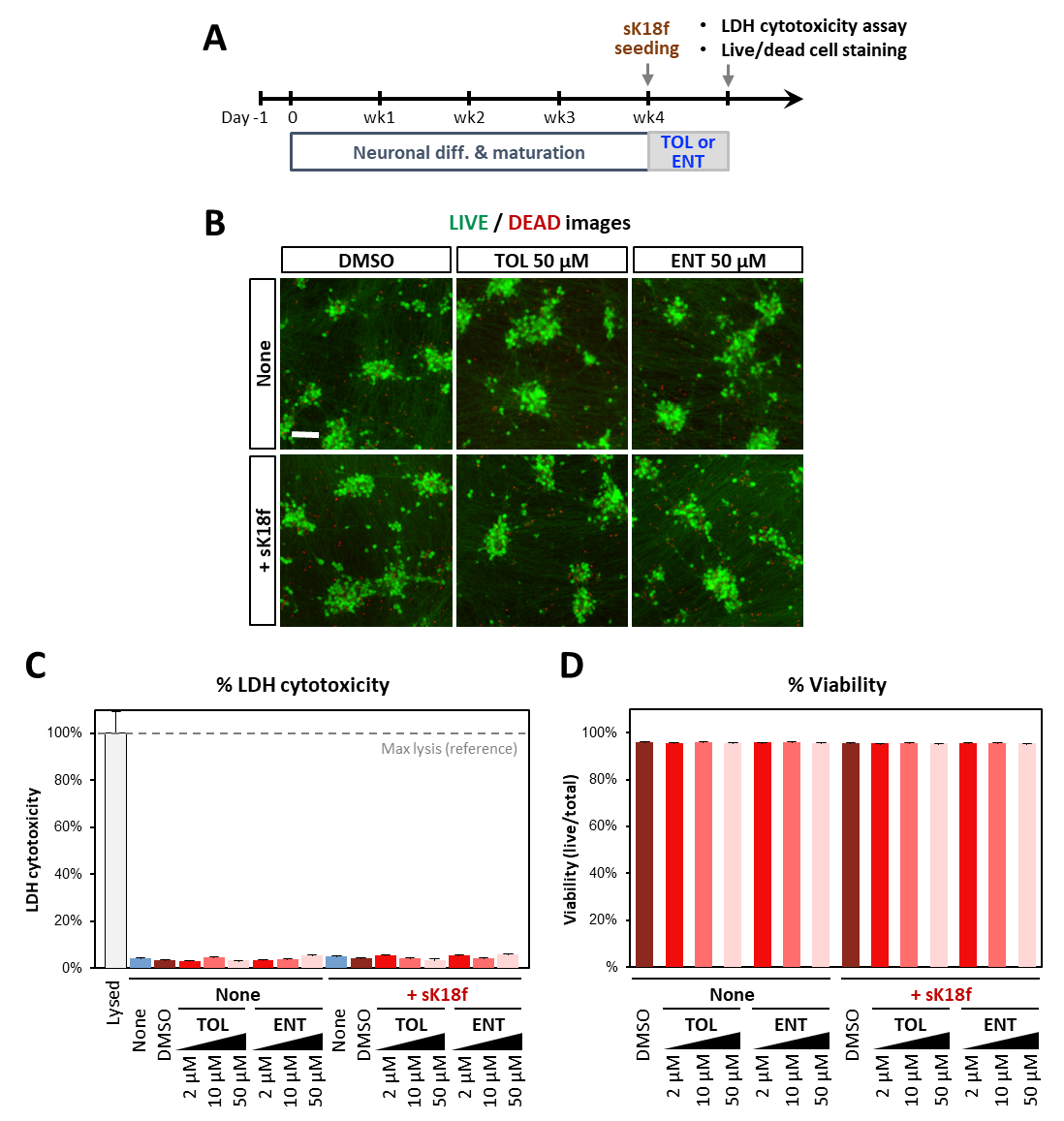

**Figure S12. TOL and ENT do not induce overt cytotoxicity in hiPSC-derived neurons.**

**(A)** Experimental timeline for toxicity assessment in NGN2-induced hiPSC-derived neurons. Neuronal differentiation and maturation were carried out until week 4, followed by sK18f seeding and treatment with TOL or ENT for five days. LIVE/DEAD staining and LDH cytotoxicity assays were performed as indicated.

**(B)** Representative LIVE/DEAD staining images of hiPSC-iNs treated with vehicle (DMSO), TOL (50 μM), or ENT (50 μM) in the absence (None) or presence (+sK18f) of sK18f. Live cells are labeled by calcein fluorescence (green) and dead cells by EthD-1 fluorescence (red). Scale bar, 100 μm.

**(C)** LDH cytotoxicity measured from culture supernatants of hiPSC-iNs treated with vehicle, TOL (2, 10, 50 μM), or ENT (2, 10, 50 μM) under None or +sK18f conditions. Maximum lysis control (reference) is shown.

**(D)** Quantification of neuronal viability from LIVE/DEAD staining for the same treatment conditions shown in panel B, reported as the percentage of viable (live) cells. Data **(C)** and **(D)** are presented as mean ± SEM from 2independent differentiations with 2-3 technical replicates per condition. Vehicle-matched DMSO was used across all conditions (final DMSO ≤ 0.25%).

### **Section B (Supplementary Tables)**

| Receptor (TAU) |  | S (E_score2)  (kcal/mol) | RMSD_refine  (Å) | E_conf | E_place | E_score1 | E_refine  (kcal/mol) |
| --- | --- | --- | --- | --- | --- | --- | --- |
| WT | Entacapone | -4.85 | 2.70 | -15.03 | -43.17 | -7.62 | -22.74 |
|  | Tolcapone | -4.82 | 3.55 | 25.20 | -46.23 | -7.03 | -25.89 |
| P301L | Entacapone | -4.80 | 2.93 | -15.97 | -35.59 | -7.50 | -23.78 |
|  | Tolcapone | -4.73 | 2.15 | 31.58 | -35.05 | -8.96 | -24.81 |
| P301S | Entacapone | -5.11 | 1.90 | -14.64 | -39.79 | -8.44 | -23.05 |
|  | Tolcapone | -4.88 | 2.30 | 31.51 | -65.79 | -7.85 | -20.66 |

**Table S1. Induced-fit docking (IFD) scores for entacapone and tolcapone against WT, P301L, and P301S tau.**

IFD was performed using MOE 2024.01 on AlphaFold2-derived tau structures, allowing flexibility of receptor side chains and backbone during ligand placement and refinement.

The table reports the following parameters:

S (E_score2), final docking score in kcal/mol (more negative values indicate stronger predicted binding);

rmsd_refine, root-mean-square deviation (Å) between the refined and initial ligand poses;

E_conf, energy associated with receptor conformational change;

E_placement, ligand placement score;

E_score1, initial rescoring energy;

E_refine, final refined binding energy in kcal/mol (more negative values indicate stronger predicted binding).

Tolcapone shows the most favorable refined binding energies for WT and P301L tau. For P301S tau, entacapone exhibits the lowest rmsd_refine value (1.90 Å), indicating a well-accommodated refined binding pose.

| Desmond | Models | Ligand | ΔG Bind | Coulomb | Covalent | H-Bond | Lipophilic | Solvation GB | Van der Waals |
| --- | --- | --- | --- | --- | --- | --- | --- | --- | --- |
| R1 | WT | ENT | -20.69 | -37.08 | 1.07 | -0.62 | -3.41 | 39.24 | -19.51 |
| R2 | WT | ENT | -20.85 | -39.40 | 1.16 | -1.01 | -3.57 | 41.30 | -18.91 |
| R3 | WT | ENT | -10.87 | -25.98 | 0.87 | -0.63 | -1.80 | 29.01 | -12.22 |
| R4 | WT | ENT | -16.83 | -28.32 | 1.10 | -0.74 | -3.20 | 31.24 | -16.77 |
| R5 | WT | ENT | -39.72 | -41.11 | 2.14 | -2.24 | -8.43 | 45.41 | -35.12 |
| R6 | WT | ENT | -21.78 | -25.76 | 1.08 | -0.65 | -5.49 | 29.14 | -19.64 |
| R7 | WT | ENT | -13.34 | -27.37 | 0.63 | -0.55 | -2.77 | 30.09 | -13.01 |
| R8 | WT | ENT | -42.33 | -49.35 | 1.99 | -1.14 | -7.37 | 48.38 | -33.86 |
| R9 | WT | ENT | -22.57 | -30.32 | 1.34 | -0.89 | -4.75 | 35.57 | -23.45 |
| R10 | WT | ENT | -33.38 | -37.78 | 1.34 | -1.32 | -7.10 | 39.86 | -27.84 |

**Table S2. MM-GBSA binding free energy decomposition (kcal/mol) for entacapone (ENT) bound to wild-type (WT) tau across 10 Desmond molecular dynamics replicas.**

Binding free energies (ΔG_bind) were calculated using the Prime MM-GBSA module on equilibrated docked poses of ENT in the WT tau binding site following 100 ns Desmond molecular dynamics simulations for each replica.

Columns report the total binding free energy (ΔG_bind) and its decomposition into energetic components: Coulomb (electrostatic interactions), Covalent, Hydrogen bond, Lipophilic, Solvation GB (generalized Born solvation term), and van der Waals contributions. More negative values indicate more favorable energetic contributions to binding.

The results show consistent interaction patterns across replicas. Two replicas (R5 and R8) display more favorable total binding energies (ΔG_bind < -39 kcal/mol), whereas the remaining replicas cluster around −20 kcal/mol, reflecting variability arising from conformational sampling during molecular dynamics simulations.

| Desmond | Models | Ligand | ΔG Bind | Coulomb | Covalent | H-Bond | Lipophilic | Solvation GB | Van der Waals |
| --- | --- | --- | --- | --- | --- | --- | --- | --- | --- |
| R1 | WT | TOL | -23.04 | -39.53 | 0.96 | -1.10 | -5.75 | 46.39 | -23.74 |
| R2 | WT | TOL | -37.80 | -37.89 | 1.35 | -1.38 | -10.30 | 43.75 | -32.26 |
| R3 | WT | TOL | -21.70 | -32.42 | 1.03 | -0.97 | -5.78 | 39.07 | -22.12 |
| R4 | WT | TOL | -20.30 | -37.80 | 0.80 | -0.82 | -4.92 | 40.69 | -17.99 |
| R5 | WT | TOL | -42.83 | -57.03 | 1.68 | -1.36 | -8.86 | 58.04 | -34.74 |
| R6 | WT | TOL | -22.54 | -29.27 | 0.91 | -0.80 | -5.60 | 33.39 | -20.09 |
| R7 | WT | TOL | -20.63 | -38.05 | 0.95 | -1.10 | -4.74 | 43.12 | -20.26 |
| R8 | WT | TOL | -24.14 | -35.00 | 0.76 | -1.52 | -6.20 | 40.10 | -21.81 |
| R9 | WT | TOL | -15.72 | -23.40 | 0.72 | -0.54 | -4.08 | 27.63 | -15.62 |
| R10 | WT | TOL | -41.53 | -33.25 | 1.31 | -0.95 | -12.94 | 39.29 | -33.17 |

**Table S3 MM‑GBSA binding free energy decomposition (kcal/mol) for tolcapone (TOL) against wild‑type (WT) tau from 10 Desmond MD replicas.**

ΔG_bind values were computed using the Prime MM‑GBSA module on equilibrated docked poses of TOL in the WT tau binding site, after 100 ns Desmond MD simulations per replica. Columns represent total binding free energy (ΔG Bind) and its decomposition: Coulomb (electrostatic), Covalent, H‑Bond, Lipophilic, Solvation GB (generalized Born solvation), and van der Waals. More negative contributions are favorable. Three high‑affinity outliers (R2, R5, R10; ΔG_bind ≤ −37.8 kcal/mol) stand out due to especially strong van der Waals and lipophilic terms, while the remaining replicas cluster around −20 to −25 kcal/mol, underscoring a reproducible yet conformationally sensitive binding mode.

| Desmond | Models | Ligand | ΔG Bind | Coulomb | Covalent | H-Bond | Lipophilic | Solvation GB | Van der Waals |
| --- | --- | --- | --- | --- | --- | --- | --- | --- | --- |
| R1 | P301S | ENT | -22.15 | -29.88 | 1.13 | -0.74 | -3.93 | 34.07 | -22.23 |
| R2 | P301S | ENT | -27.83 | -42.73 | 1.33 | -1.21 | -4.39 | 45.20 | -25.06 |
| R3 | P301S | ENT | -25.36 | -39.98 | 1.23 | -0.96 | -5.55 | 45.15 | -25.04 |
| R4 | P301S | ENT | -19.62 | -34.83 | 0.93 | -0.78 | -3.77 | 37.47 | -18.29 |
| R5 | P301S | ENT | -39.06 | -55.28 | 1.82 | -1.84 | -6.94 | 54.72 | -31.36 |
| R6 | P301S | ENT | -28.43 | -58.97 | 1.43 | -1.12 | -3.06 | 63.13 | -29.83 |
| R7 | P301S | ENT | -29.64 | -44.17 | 1.55 | -1.60 | -5.07 | 47.66 | -26.91 |
| R8 | P301S | ENT | -47.32 | -59.92 | 1.77 | -2.42 | -6.95 | 58.16 | -36.71 |
| R9 | P301S | ENT | -23.67 | -33.02 | 1.29 | -1.09 | -4.45 | 35.40 | -21.22 |
| R10 | P301S | ENT | -24.89 | -40.02 | 1.23 | -1.20 | -3.83 | 45.41 | -26.08 |

**Table S4. MM-GBSA binding free energy decomposition (kcal/mol) for entacapone (ENT) bound to the P301S tau mutant across 10 Desmond molecular dynamics replicas.**

Binding free energies (ΔG_bind) were calculated using the Prime MM-GBSA module on equilibrated docked poses of ENT in the P301S tau binding site following 100 ns Desmond molecular dynamics simulations for each replica.

Columns report the total binding free energy (ΔG_bind) and its decomposition into energetic components: Coulomb (electrostatic interactions), Covalent, Hydrogen bond, Lipophilic, Solvation GB (generalized Born solvation term), and van der Waals contributions. More negative values indicate more favorable energetic contributions.

Individual replicas exhibit variability consistent with conformational sampling during molecular dynamics simulations, with some showing more favorable total binding energies (e.g., R8: −47.32 kcal/mol) and stronger Coulomb and van der Waals contributions.

| Desmond | Models | Ligand | ΔG Bind | Coulomb | Covalent | H-Bond | Lipophilic | Solvation GB | Van der Waals |
| --- | --- | --- | --- | --- | --- | --- | --- | --- | --- |
| R1 | P301S | TOL | -19.27 | -38.27 | 0.95 | -0.98 | -2.50 | 41.53 | -19.66 |
| R2 | P301S | TOL | -16.77 | -35.08 | 0.66 | -0.38 | -4.21 | 38.36 | -15.96 |
| R3 | P301S | TOL | -39.12 | -51.63 | 1.07 | -1.36 | -8.49 | 54.33 | -32.63 |
| R4 | P301S | TOL | -34.37 | -47.66 | 1.36 | -2.24 | -6.66 | 54.72 | -32.57 |
| R5 | P301S | TOL | -26.52 | -39.79 | 1.16 | -0.91 | -6.82 | 44.87 | -24.54 |
| R6 | P301S | TOL | -28.38 | -43.79 | 1.10 | -1.34 | -5.85 | 50.01 | -27.26 |
| R7 | P301S | TOL | -34.42 | -63.85 | 0.97 | -1.88 | -3.64 | 67.14 | -31.02 |
| R8 | P301S | TOL | -30.73 | -49.12 | 1.11 | -1.06 | -6.99 | 54.55 | -27.85 |
| R9 | P301S | TOL | -20.98 | -37.66 | 1.01 | -0.86 | -4.83 | 42.88 | -20.61 |
| R10 | P301S | TOL | -20.41 | -40.42 | 1.28 | -1.04 | -3.92 | 44.00 | -19.52 |

**Table S5. MM-GBSA binding free energy decomposition (kcal/mol) for tolcapone (TOL) bound to the P301S tau mutant across 10 Desmond molecular dynamics replicas.**

Binding free energies (ΔG_bind) were calculated using the Prime MM-GBSA module on equilibrated docked poses of TOL in the P301S tau binding site following 100 ns Desmond molecular dynamics simulations for each replica.

Columns report the total binding free energy (ΔG_bind) and its decomposition into energetic components: Coulomb (electrostatic interactions), Covalent, Hydrogen bond, Lipophilic, Solvation GB (generalized Born solvation term), and van der Waals contributions. More negative values indicate more favorable energetic contributions.

The calculated binding energies vary across replicas, consistent with conformational sampling during molecular dynamics simulations. Some replicas exhibit more favorable total binding energies (e.g., R3: −39.12 kcal/mol), reflecting stronger Coulomb and van der Waals contributions.

| Desmond | Models | Ligand | ΔG Bind | Coulomb | Covalent | H-Bond | Lipophilic | Solvation GB | Van der Waals |
| --- | --- | --- | --- | --- | --- | --- | --- | --- | --- |
| R1 | P301L | ENT | -36.64 | -45.83 | 2.18 | -1.67 | -6.98 | 51.47 | -35.15 |
| R2 | P301L | ENT | -35.11 | -49.15 | 1.32 | -1.03 | -6.83 | 53.48 | -32.39 |
| R3 | P301L | ENT | -16.97 | -28.13 | 1.12 | -0.78 | -3.28 | 31.72 | -17.43 |
| R4 | P301L | ENT | -48.44 | -45.79 | 2.21 | -2.27 | -9.21 | 49.62 | -42.27 |
| R5 | P301L | ENT | -17.94 | -29.40 | 0.74 | -0.82 | -4.38 | 30.59 | -14.62 |
| R6 | P301L | ENT | -35.61 | -56.80 | 1.74 | -1.46 | -6.52 | 59.90 | -32.42 |
| R7 | P301L | ENT | -13.44 | -24.81 | 0.85 | -0.64 | -2.60 | 27.98 | -14.00 |
| R8 | P301L | ENT | -22.21 | -30.51 | 1.07 | -0.78 | -5.03 | 34.68 | -21.33 |
| R9 | P301L | ENT | -44.46 | -45.13 | 1.21 | -2.31 | -9.73 | 48.23 | -36.54 |
| R10 | P301L | ENT | -7.09 | -18.85 | 0.45 | -0.34 | -1.42 | 20.19 | -6.99 |

**Table S6. MM-GBSA binding free energy decomposition (kcal/mol) for entacapone (ENT) bound to the P301L tau mutant across 10 Desmond molecular dynamics replicas.**

Binding free energies (ΔG_bind) were calculated using the Prime MM-GBSA module on equilibrated docked poses of ENT in the mutant (P301L) tau binding site following 100 ns Desmond molecular dynamics simulations for each replica.

Columns report the total binding free energy (ΔG_bind) and its decomposition into energetic components: Coulomb (electrostatic interactions), Covalent, Hydrogen bond, Lipophilic, Solvation GB (generalized Born solvation term), and van der Waals contributions. More negative values indicate more favorable energetic contributions.

Compared to WT, P301L exhibits more frequently high-affinity replicas, with standout values driven by intensified Coulomb and van der Waals interactions (e.g., R4: −48.44 kcal/mol).

| Desmond | Models | Ligand | ΔG Bind | Coulomb | Covalent | H-Bond | Lipophilic | Solvation GB | Van der Waals |
| --- | --- | --- | --- | --- | --- | --- | --- | --- | --- |
| R1 | P301L | TOL | -34.23 | -44.42 | 1.45 | -1.43 | -6.38 | 49.45 | -31.84 |
| R2 | P301L | TOL | -23.71 | -46.08 | 1.18 | -1.04 | -4.96 | 51.44 | -23.71 |
| R3 | P301L | TOL | -18.78 | -31.37 | 0.89 | -1.02 | -4.76 | 35.19 | -17.36 |
| R4 | P301L | TOL | -5.92 | -16.09 | 0.28 | -0.21 | -1.71 | 17.41 | -5.50 |
| R5 | P301L | TOL | -35.41 | -48.25 | 1.41 | -1.40 | -9.72 | 54.45 | -31.15 |
| R6 | P301L | TOL | -36.71 | -39.13 | 0.94 | -1.41 | -8.49 | 43.00 | -30.13 |
| R7 | P301L | TOL | -37.40 | -57.21 | 1.46 | -1.59 | -7.91 | 61.59 | -33.63 |
| R8 | P301L | TOL | -22.33 | -36.82 | 1.01 | -1.01 | -5.64 | 42.32 | -21.56 |
| R9 | P301L | TOL | -35.82 | -51.47 | 1.08 | -1.07 | -10.60 | 59.25 | -32.16 |
| R10 | P301L | TOL | -38.14 | -43.13 | 1.66 | -1.45 | -10.48 | 49.41 | -32.82 |

**Table S7. MM-GBSA binding free energy decomposition (kcal/mol) for tolcapone (TOL) bound to the P301L tau mutant across 10 Desmond molecular dynamics replicas.**

Binding free energies (ΔG_bind) were calculated using the Prime MM-GBSA module on equilibrated docked poses of TOL in the P301L tau binding site following 100 ns Desmond molecular dynamics simulations for each replica.

Columns report the total binding free energy (ΔG_bind) and its decomposition into energetic components: Coulomb (electrostatic interactions), Covalent, Hydrogen bond, Lipophilic, Solvation GB (generalized Born solvation term), and van der Waals contributions. More negative values indicate more favorable energetic contributions.

Several replicas show relatively favorable total binding energies (e.g., R7: −37.40 kcal/mol), with notable contributions from Coulomb and van der Waals terms. Overall variability across replicas reflects conformational sampling during molecular dynamics simulations.

### **Section C (Supplementary Methods)**

***SH-SY5Y* *Cell Viability Assay***

Cytotoxicity of tolcapone (TOL) and entacapone (ENT) was assessed using a colorimetric MTS viability assay (CellTiter 96® AQueous One Solution, Promega) according to the manufacturer’s instructions and an established in-house protocol. SH-SY5Y cells were seeded at 10,000 cells per well in 96-well plates and treated with increasing concentrations of TOL or ENT for 72 h. DMSO-treated cells served as vehicle controls, with a final DMSO concentration ≤0.125% (v/v).

After treatment, MTS reagent was added, and absorbance was measured using a multimode microplate reader. Background-subtracted absorbance values were normalized to vehicle controls and expressed as percentage viability. Concentration–response relationships were analyzed by nonlinear regression using a four-parameter logistic model in GraphPad Prism. Raw absorbance values and full regression outputs are available via Zenodo (see Data Availability Statement).

***Homology Modeling***

WT Tau, P301L, and P301S Tau sequences were modeled using AlphaFold2 [1], which predicts three-dimensional protein structures from amino acid sequences using multiple sequence alignments (MSAs) and structural templates (Fig. S5A–C). MSAs were generated using UniRef and MGnify databases with MMseqs2/HHblits, following standard AlphaFold2 protocols [2].

Model quality was evaluated using predicted Local Distance Difference Test (pLDDT) scores and Predicted Aligned Error (PAE) to assess residue-level confidence (Fig. S6A–C and Fig. S7A–C). Sequence coverage plots were used to assess MSA depth and alignment quality (Fig. S8A–C). The highest-ranked model (rank_0) with the best predicted confidence metrics was selected for each construct (WT: pLDDT 50.8, pTM 0.116; P301L: pLDDT 51.8, pTM 0.112; P301S: pLDDT 52.0, pTM 0.155). Structural validation was further assessed using Ramachandran plots (Fig. S9A–C).

***Molecular docking***

AlphaFold2-derived models of WT Tau, P301L, and P301S were prepared in MOE 2024.01. Structures were protonated using Protonate3D at pH 7.4 and energy-minimized with a rigid backbone. Binding pockets were identified using Site Finder, and a single-pocket dummy model was generated for docking.

Ligands (ENT and TOL) were prepared using LigPrep to generate three-dimensional conformations, protonation states, and stereoisomers.

Initial docking was performed using a rigid-receptor approach with Triangle Matcher (30 poses), followed by force field refinement. Induced fit docking was then applied to optimize the top 10 poses with side-chain flexibility. The best pose was selected based on Gibbs free binding energy (ΔG binding), RMSD, and binding affinity [3]. Reported output parameters include final score (S), RMSD, conformer energy (E_conf), and intermediate docking stage scores. Lower S scores indicate more favorable predicted binding. Detailed results are provided in Table S1.

***Molecular dynamics simulations and binding free energy calculations***

Molecular dynamics (MD) simulations were performed using Desmond in Schrödinger Maestro 2025-1 to evaluate ligand–protein complex stability [4]. Systems were solvated in an SP3 water box, neutralized with Na⁺ and Cl⁻ ions, and equilibrated prior to production runs. Simulations were conducted for 100 ns at 300 K and 1 atm using the OPLS4 force field.

Ligand–protein interactions were analyzed using Simulation Interaction Diagram (SID) analysis, categorizing contacts into hydrogen bonds, hydrophobic interactions, ionic interactions, and water bridges. To improve statistical robustness, 10 independent MD simulations were performed for each complex.

Binding free energy (ΔG_bind) was calculated using the MM/GBSA method implemented in Schrödinger Maestro 2025-1 [5]. This approach accounts for van der Waals, electrostatic, solvation, and entropic contributions. Binding energy values for WT, P301L, and P301S complexes are summarized in Tables S2–S7.

**Per-Residue Decomposition**

Per-residue energy decomposition was performed using the MM-GBSA implementation in Desmond (Schrödinger Suite) to quantify the contribution of individual residues to total binding free energy (ΔG_bind). Energy contributions were partitioned into van der Waals, electrostatic, solvation, and entropic components.

Following MD simulations, representative trajectory snapshots (e.g., 100–500 frames across 10 replicates) were analyzed using Prime MM-GBSA decomposition according to:

ΔG_res = ΔE_vdW + ΔE_ele + ΔG_solv − TΔS

Residues with ΔG_res < −3 kcal/mol were defined as stabilizing interaction hotspots. Source data for per-residue decomposition is available via Zenodo (see Data Availability Statement).

### **Section D (Uncropped Western Blots Images)**

**Uncropped blot images - Figure 3 — Triton-X-100- insoluble blots of cells transduced with fibrils of wildtype R2R3 tau constructs formed in the absence (CTL) or presence of tolcapone (TOL) or entacapone (ENT)**

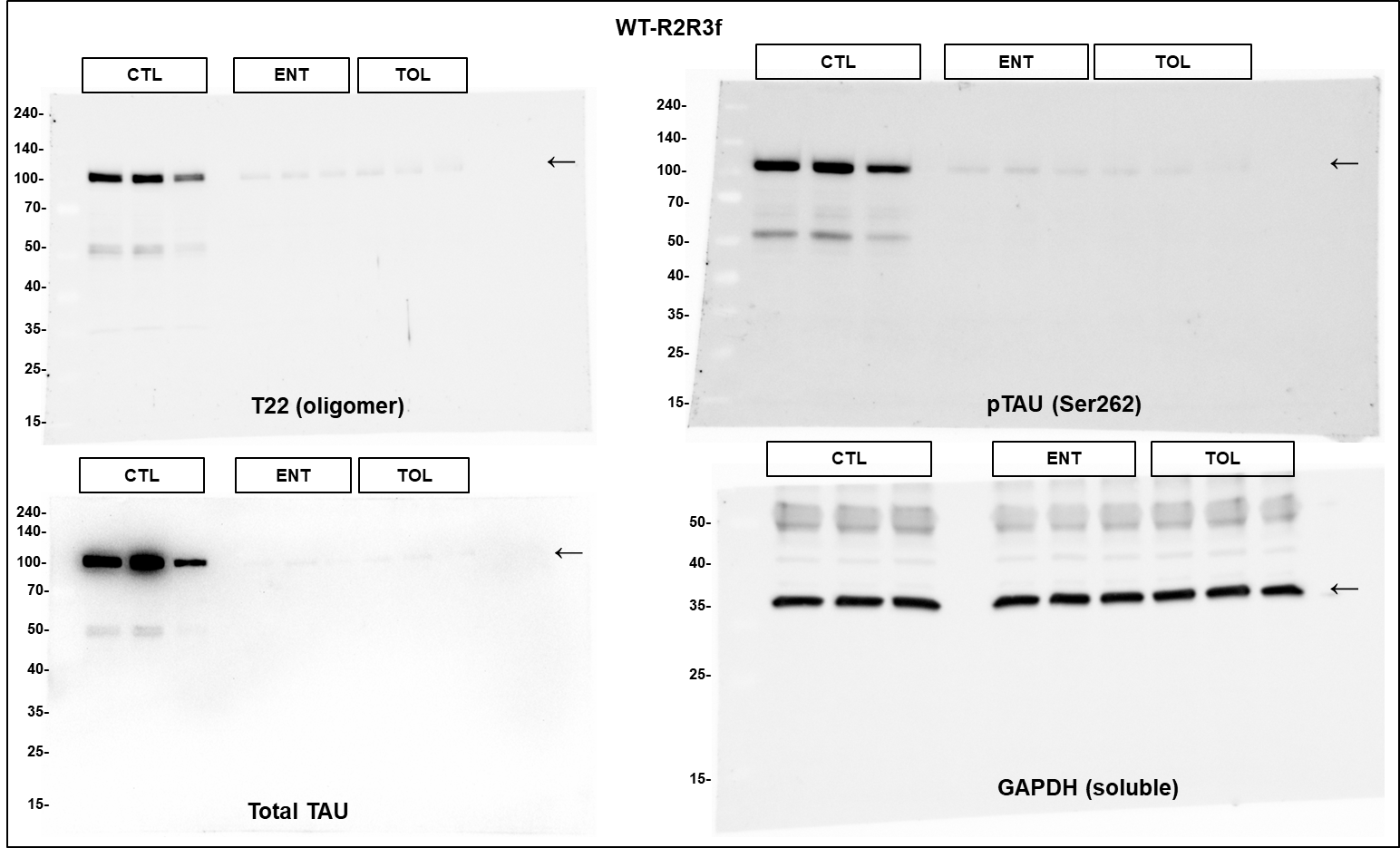

**Uncropped blot images - Figure 3 — Triton-X-100- insoluble blots of cells transduced with fibrils of mutant P301S R2R3 tau constructs formed in the absence (CTL) or presence of tolcapone (TOL) or entacapone (ENT)**

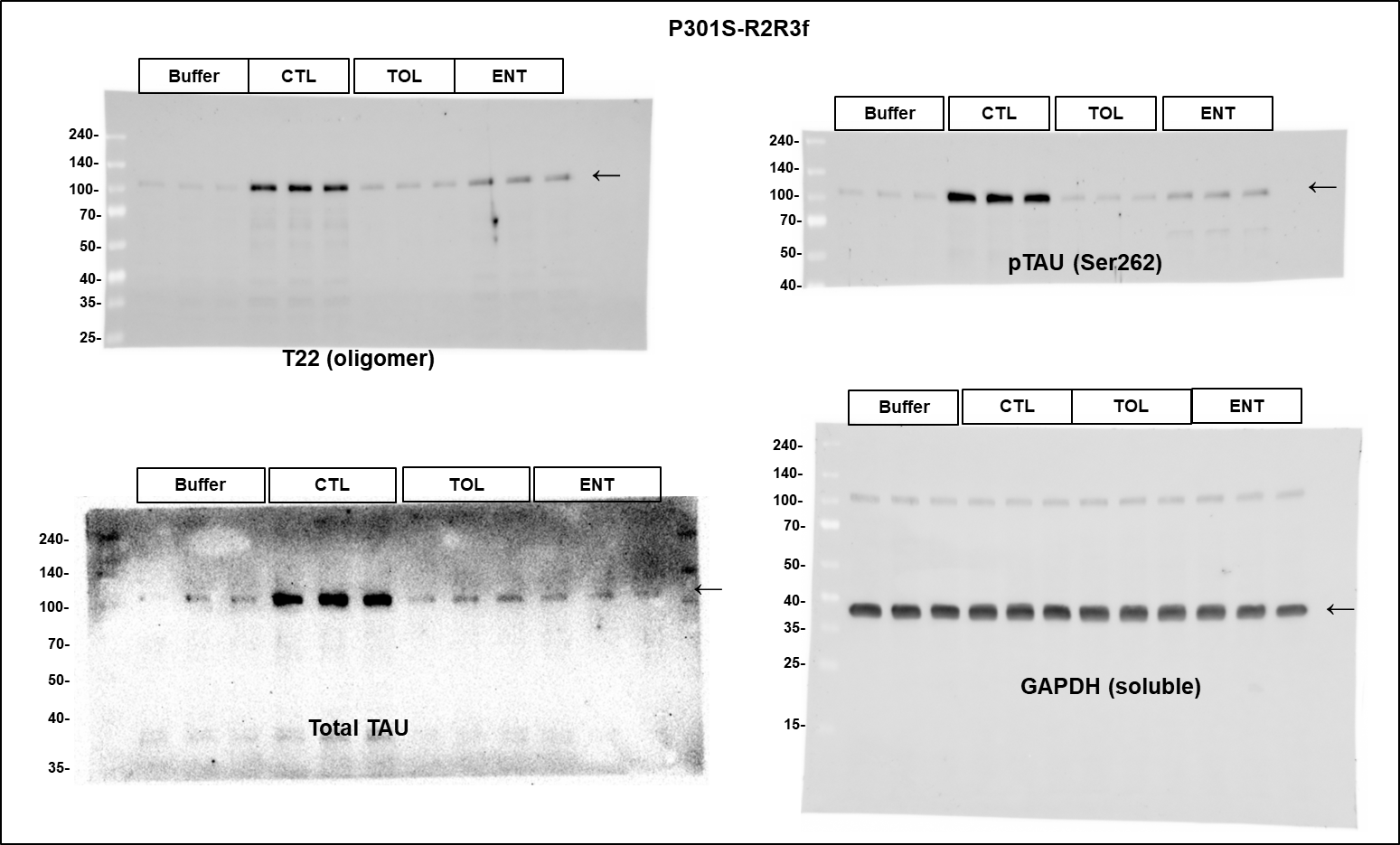

**Figure 5 — Sarkosyl insoluble tau**

| 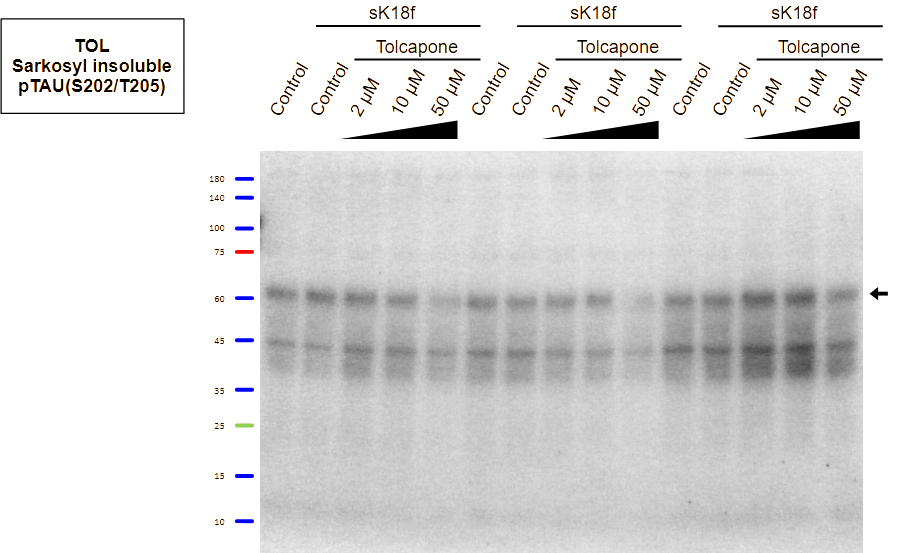 | 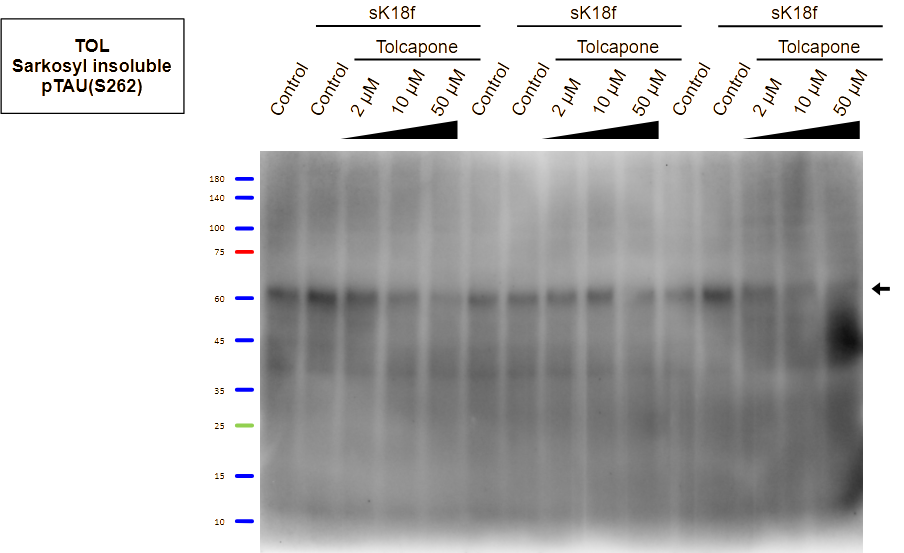 |
| --- | --- |
| 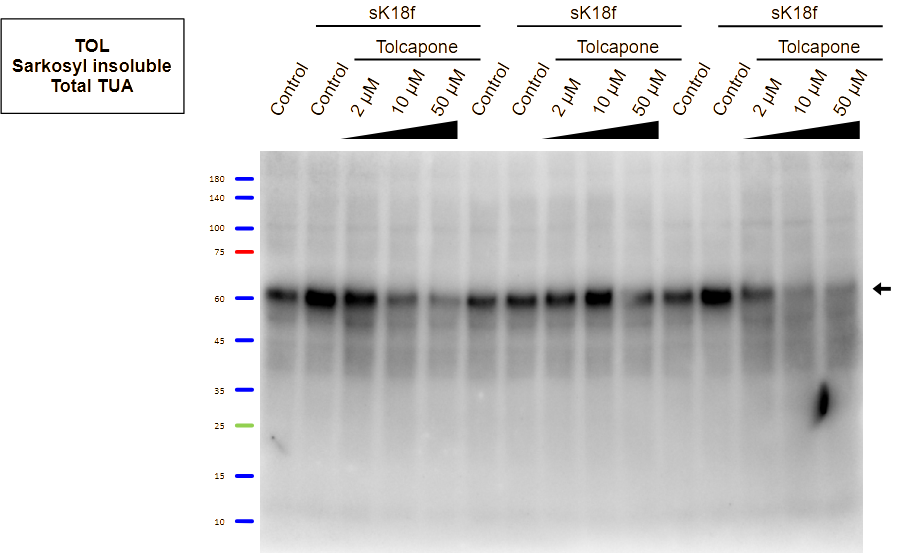 | 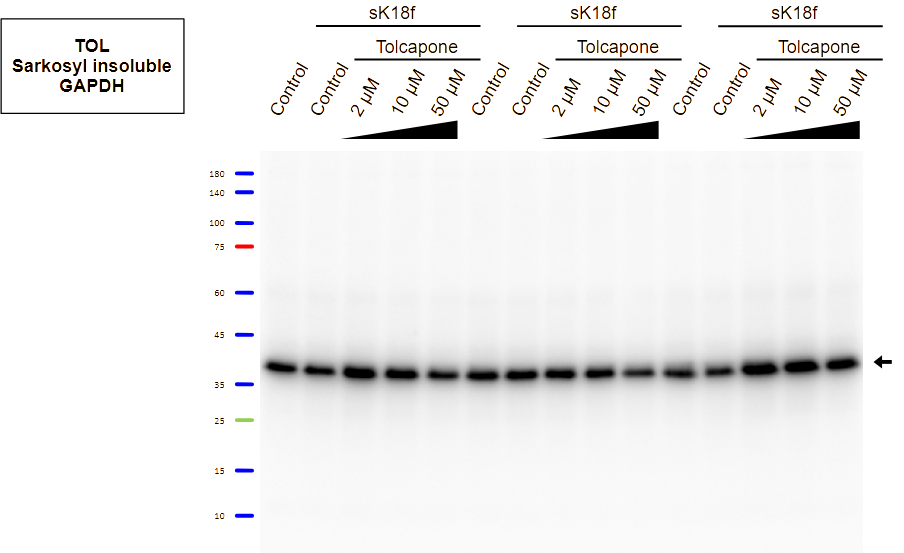 |

| 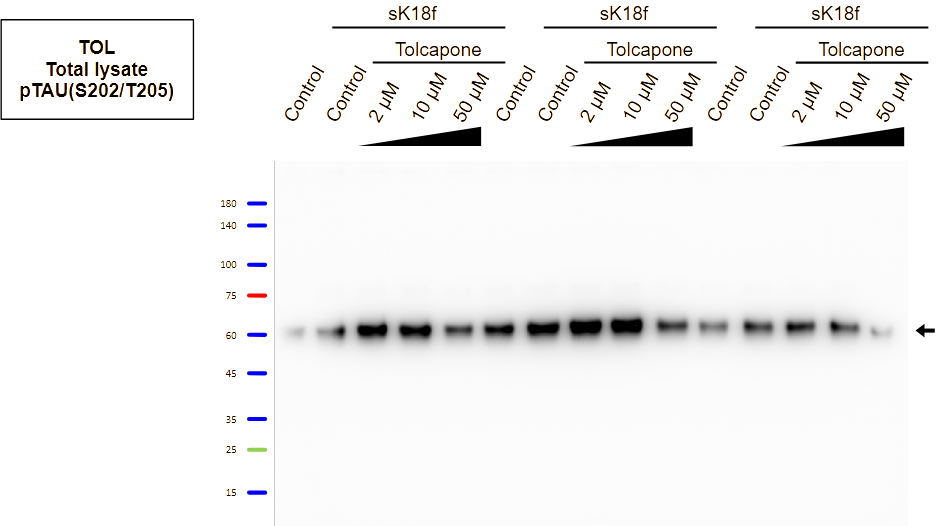 | 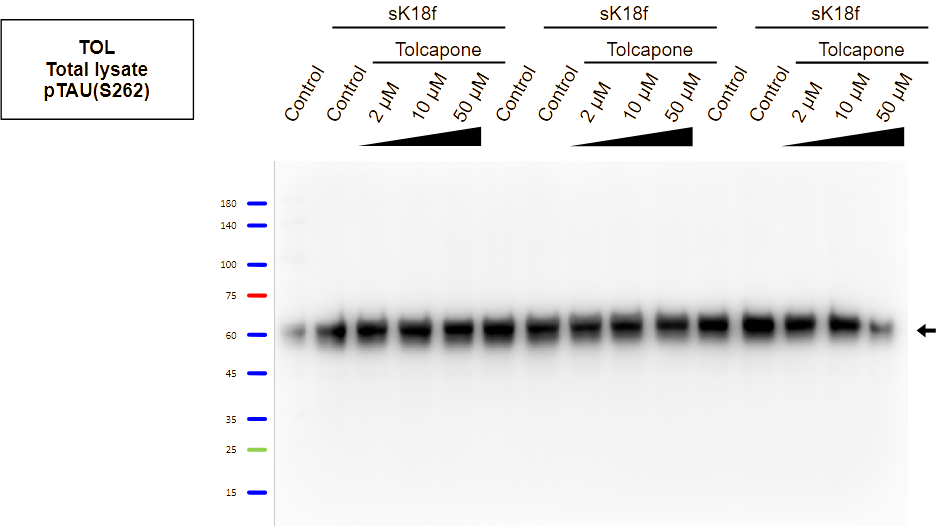 |
| --- | --- |
| 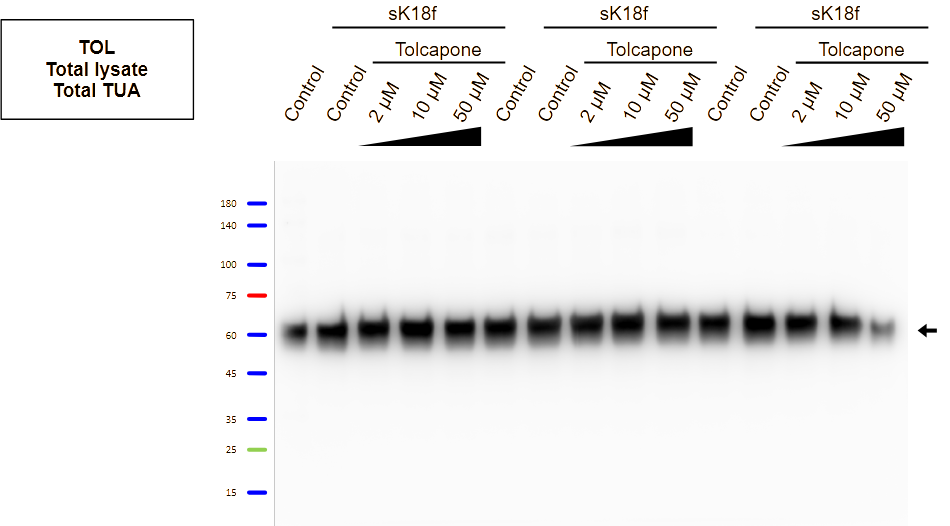 | 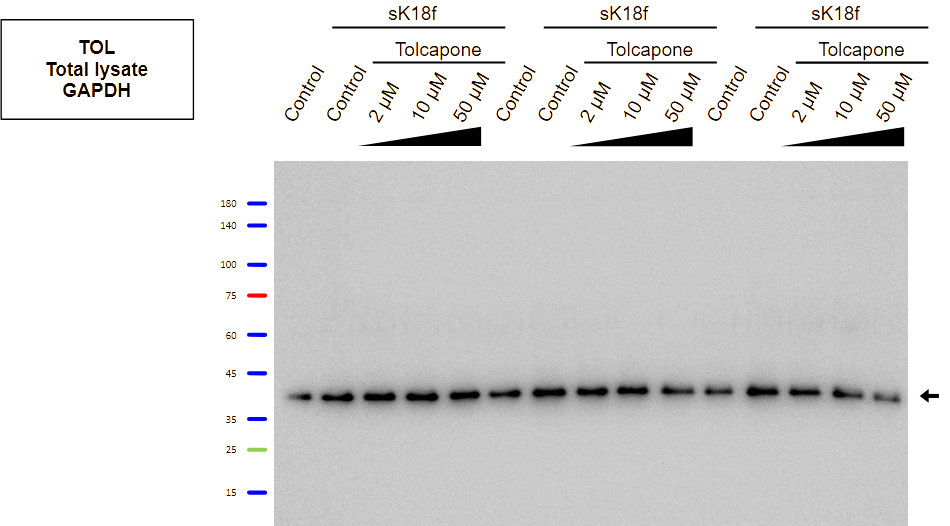 |
| 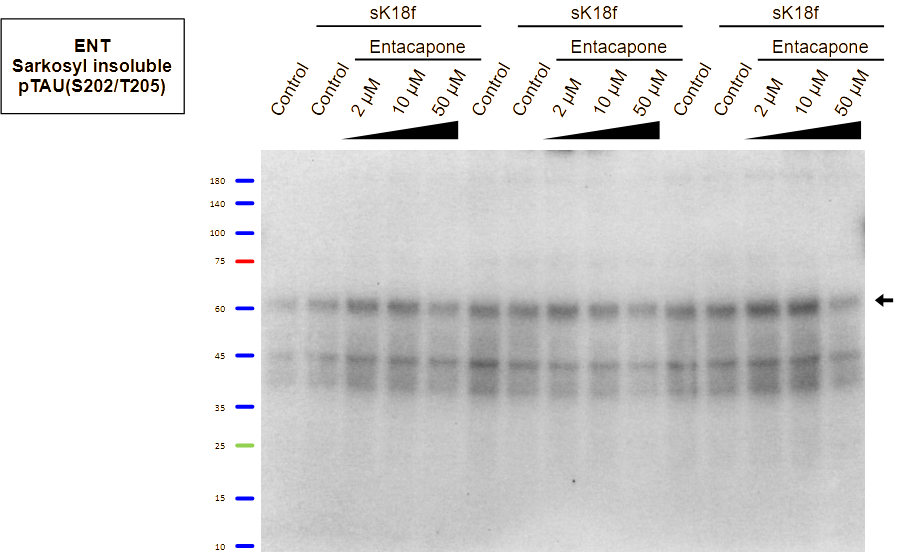 | 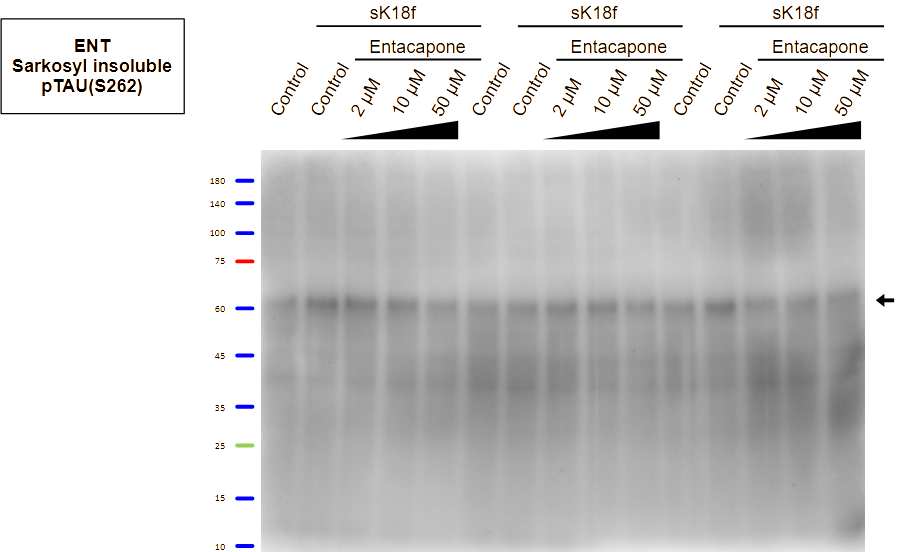 |
| 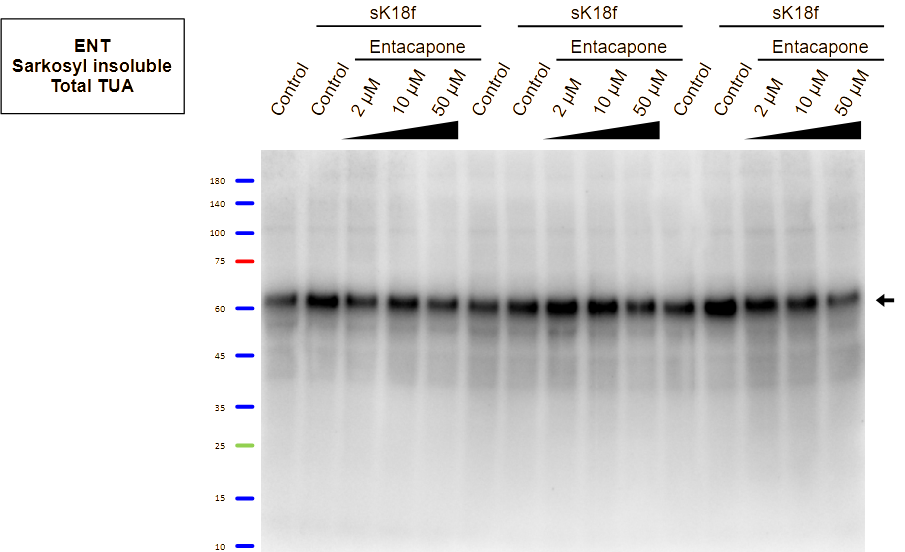 | 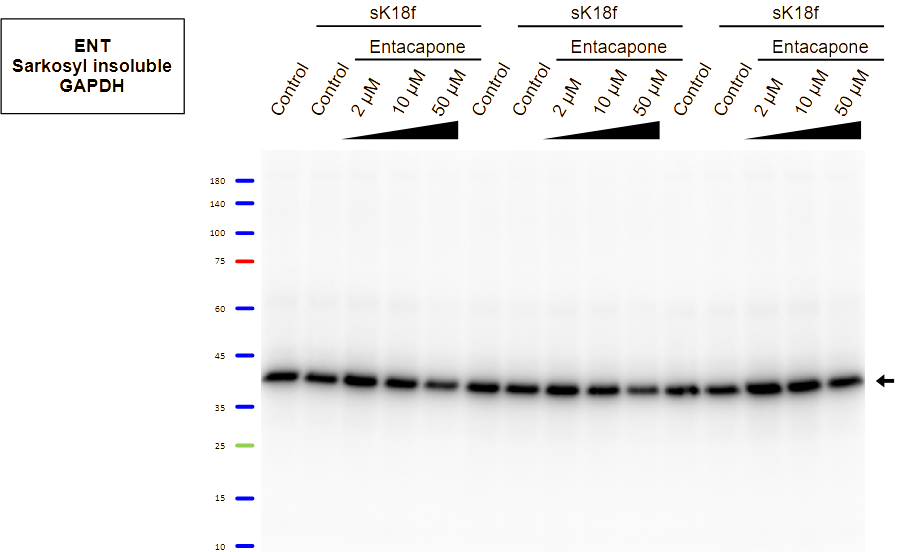 |
| 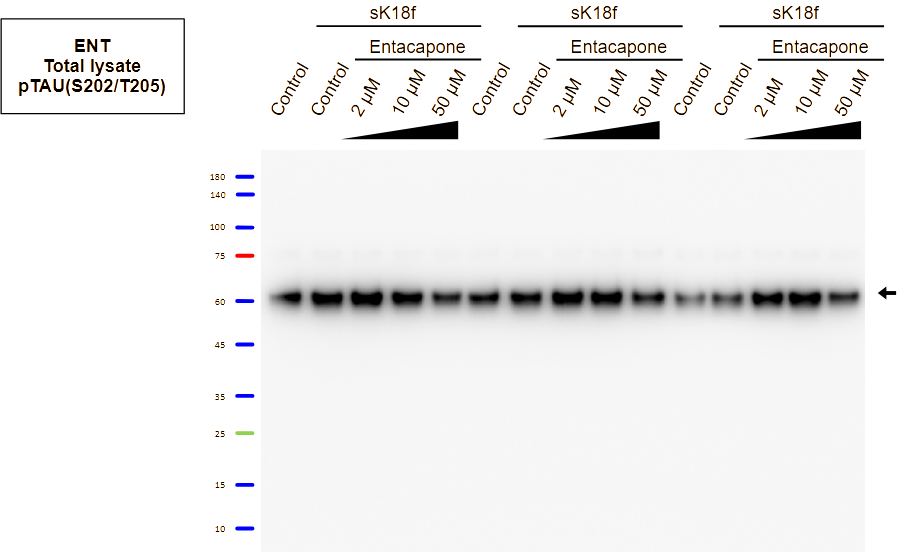 | 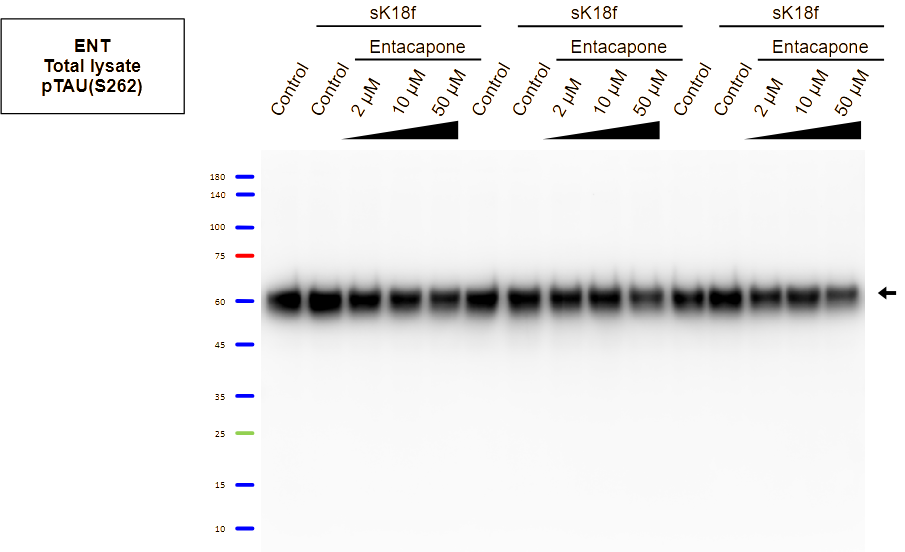 |
| 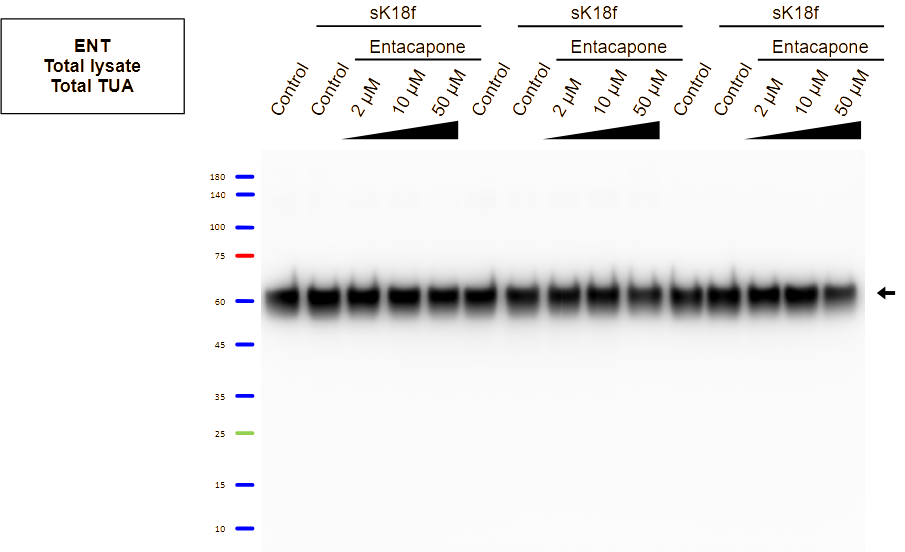 | 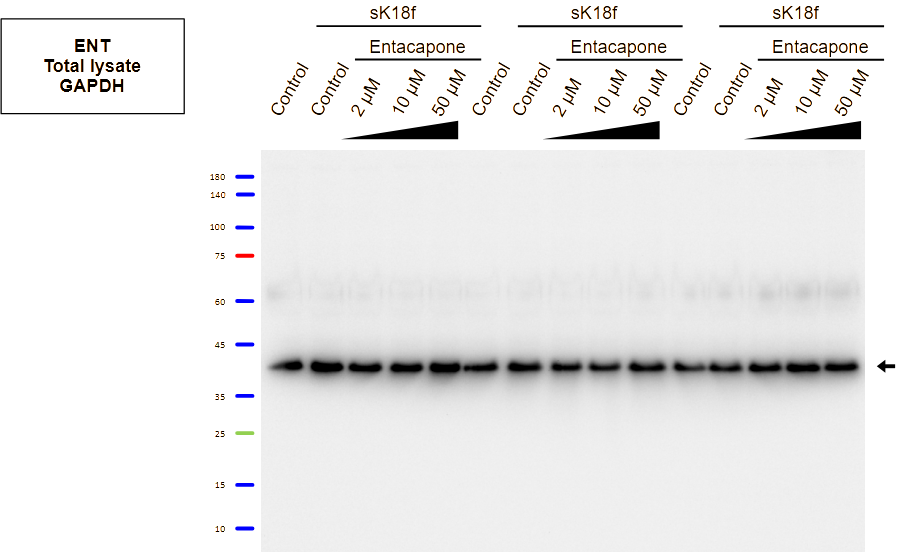 |
